# Filament formation activates protease and ring nuclease activities of CRISPR SAVED-Lon

**DOI:** 10.1101/2024.05.08.593097

**Authors:** Dalia Smalakyte, Audrone Ruksenaite, Giedrius Sasnauskas, Giedre Tamulaitiene, Gintautas Tamulaitis

## Abstract

To combat phage infection, type III CRISPR-Cas systems utilize cyclic oligoadenylates (cA_n_) signaling to activate various auxiliary effectors, including the CRISPR-associated SAVED-Lon protease CalpL, which forms a tripartite effector system together with an anti-σ factor, CalpT, and an ECF-like σ factor, CalpS. Here we report the characterization of the *Candidatus* Cloacimonas acidaminovorans CalpL-CalpT-CalpS. We demonstrate that cA_4_ binding triggers CalpL filament formation and activates it to cleave CalpT within the CalpT-CalpS dimer. This cleavage exposes the CalpT C-degron, which targets it for further degradation by cellular proteases. Consequently, CalpS is released to bind to RNA polymerase, causing growth arrest in *E. coli*. Furthermore, the CalpL-CalpT-CalpS system is regulated by the SAVED domain of CalpL, which is a ring nuclease that cleaves cA_4_ in a sequential three-step mechanism. These findings provide key mechanistic details for the activation, proteolytic events, and regulation of the signaling cascade in the type III CRISPR-Cas immunity.

## INTRODUCTION

CRISPR-Cas systems are adaptive immune systems that protect prokaryotes against invasive mobile genetic elements^1^. While all CRISPR-Cas types use a crRNA-guided protein or protein complex to target and degrade foreign nucleic acids that are complementary to the guide molecule, type III CRISPR-Cas systems use an additional layer of defense, a cyclic oligoadenylate (cA_n,_ n=3-6)-based signaling pathway to activate accessory effector proteins^2^. Upon binding to complementary foreign RNA, Csm/Cmr interference complex activates the Palm domain in the large subunit Cas10 of the complex to synthesize cA_n_ from ATP^3,4^. cA_n_ are subsequently bound by accessory proteins and this binding triggers the enzymatic activity of the fused effector domains. The most common domain architecture of the accessory proteins associated with the type III-A/B/D systems is a sensor CARF (CRISPR-Cas Associated Rossmann Fold) domain fused to various effector domains^5^. In recent years, several of CARF auxiliary type III effectors have been characterized including RNA nucleases (Csm6 and Csx1)^3,6^, DNA nucleases (Can1, Can2, Card1)^7–9^, transcriptional regulator^10^ and translational inhibitor^11^, CARF-transmembrane helix fusion^12,13^. Moreover, an association between type III CRISPR-Cas systems and effector proteins containing sensory SAVED domains (SMODS-Associated and fused to Various Effector Domains) instead of the CARF domain has also been observed^5,14^. SAVED domains are closely associated with another prokaryotic defense system named CBASS (cyclic oligonucleotide-based antiphage signaling system), which also depends on cyclic nucleotide signaling and effector protein activation^15^. A recent study by Rouillon et al. has identified the SAVED domain and Lon protease fusion protein CalpL as an associated effector in the type III-B CRISPR-Cas system of *Sulfurihydrogenibium spp. YO3AOP1* (SsCalpL)^16^. In bacteria stand-alone Lon proteases are responsible for the degradation of misfolded proteins, cleaving them with little sequence specificity^17^, while SsCalpL selectively binds cA_4_ and specifically cleaves CalpT protein within the stable heterocomplex of CalpT-CalpS. The crystal structure of the CalpL+cA_4_ complex showed the monomeric state of the protein. In contrast, dynamic light scattering and SAXS experiments revealed the dimerization of CalpL in a cA_4_-dependent and protein concentration-dependent manner. Therefore, the cleavage of CalpT by activated CalpL *in trans* was proposed, however, the detailed mechanism of CalpL activation and CalpT cleavage remains to be elucidated.

In contrast to type III-A/B/D systems, type III-E systems lack a large subunit Cas10 and thus cannot synthesize cA_n_. Instead, these systems depend on protein-protein interactions to activate auxiliary effector. Recently, it has been demonstrated that the type III-E interference complex directly interacts with TRP-CHAT protease^18,19^. The TRP-CHAT protease activity is triggered by the binding of the target RNA to the III-E complex^19^. Interestingly, both TRP-CHAT and CalpL appear to be components of a tripartite effector system consisting of a protease, an anti-σ factor and a σ factor. The activated TRP-CHAT protease in the III-E system has been shown to specifically cleave Csx30 protein in the Csx30-CASP-σ pair and this cleavage releases CASP-σ, a σ factor, for further control of gene expression^19^. Similarly, cA_4_-activated CalpL cleaves the CalpT protein in the CalpT-CalpS protein pair. CalpS expressed individually has been shown to copurify with RNAP, suggesting that the CalpT-CalpS pair also functions as an anti-σ/σ factor pair^16,20^. However, a CalpT cleavage product, CalpT_23_, remains bound to CalpS in a stable complex, raising the question of how CalpS is released to interact with RNAP.

The activation of TRP-CHAT is controlled by RNA binding, and its activity is inhibited, when the target RNA, bound by the interference complex is cleaved^21^. In the cA_n_-based signaling pathway, cA_n_ production is also halted once the target RNA is cleaved by the Csm/Cmr complex^3,22^. However, if synthesized cA_n_ are not degraded, they can continue to stimulate auxiliary effector proteins. Hence, it is crucial to regulate the signaling pathway to attenuate the response and prevent excessive activation of effector proteins, which could lead to cell death. The scenario of abortive infection is observed in CBASS systems, where no component responsible for degradation of secondary messengers has been identified^23^. However, type III CRISPR-Cas systems have additional enzymatic activity to cleave the messenger molecules to inactivate the auxiliary effector^2^. This can involve specialized sole proteins called CRISPR ring nucleases (Crn)^24,25^ or the CARF domain of the effector itself detects cA_n_ signal and cleaves the bound cA_n_ molecule to modulate signaling^26–29^. Interestingly, Crn2 family proteins can be found fused to effector Csx1, the CARF domain of which lacks ring nuclease activity^30^. However, the regulation mechanism of the CalpL-CalpT-CalpS tripartite effector system remains to be resolved.

Here, we focus on the molecular mechanism of the type III-A CRISPR-Cas-associated CalpL-CalpT-CalpS effector system from *Candidatus* Cloacimonas acidaminovorans strain Evry (CCa). We demonstrate that the CCaCalpL effector, a fusion protein of the cA_n_-sensor SAVED and Lon protease domains, functions as a cA_4_-triggered protease. It specifically cleaves the CCaCalpT anti-σ factor in the CCaCalpT-CalpS anti-σ/σ factor heterodimer complex. Additionally, we show that CCaCalpL acts as a self-limiting effector, with the SAVED domain specifically cleaving its activator cA_4_. We also present the cryo-EM structure of the activator-induced filament of CCaCalpL, highlighting the critical role of filament formation for the protease and ring nuclease activities. Lastly, we show that following CCaCalpL cleavage, the C-degron sequence is exposed on the CCaCalpT_24N_ fragment, targeting it for subsequent degradation by cellular proteases. This degradation step is essential for the release of the CCaCalpS σ factor from the heterodimer complex which then binds RNA polymerase and induces growth arrest in *E. coli*.

## RESULTS

### *Candidatus* C. acidaminovorans type III-A CRISPR-Cas encodes cA_4_-activated CalpL-CalpT-CalpS cascade

The type III-A CRISPR-Cas system of *Ca*. C. acidaminovorans consists of *csm1*-5 genes encoding the Csm interference complex (CCaCsm), *cas6* encoding the crRNA processing protein, the adaptation genes *cas1* and *cas2*, and the CRISPR array (Figure 1A). In addition to auxiliary effector *cami1* ^11^, this locus contains several adjacent genes (Figure 1A). According to HHpred analysis, CLOAM_RS04055 is a CARF protein with an additional C-terminal adenosine deaminase domain, whereas the next three genes are homologous to the *Sulfurihydrogenibium* spp. YO3AOP1 CalpL, CalpT and CalpS proteins respectively^16^: CLOAM_RS04060 (henceforth - CCaCalpL) comprises an N-terminal domain, a middle Lon protease domain and a C-terminal SAVED domain (Figure 1B), CLOAM_RS04065 (CCaCalpT) consists of a MazF-like domain and a DUF2080 domain, and CLOAM_RS04070 (CCaCalpS) is homologous to the extracytoplasmic function (ECF)-like σ factor.

**Figure 1.**
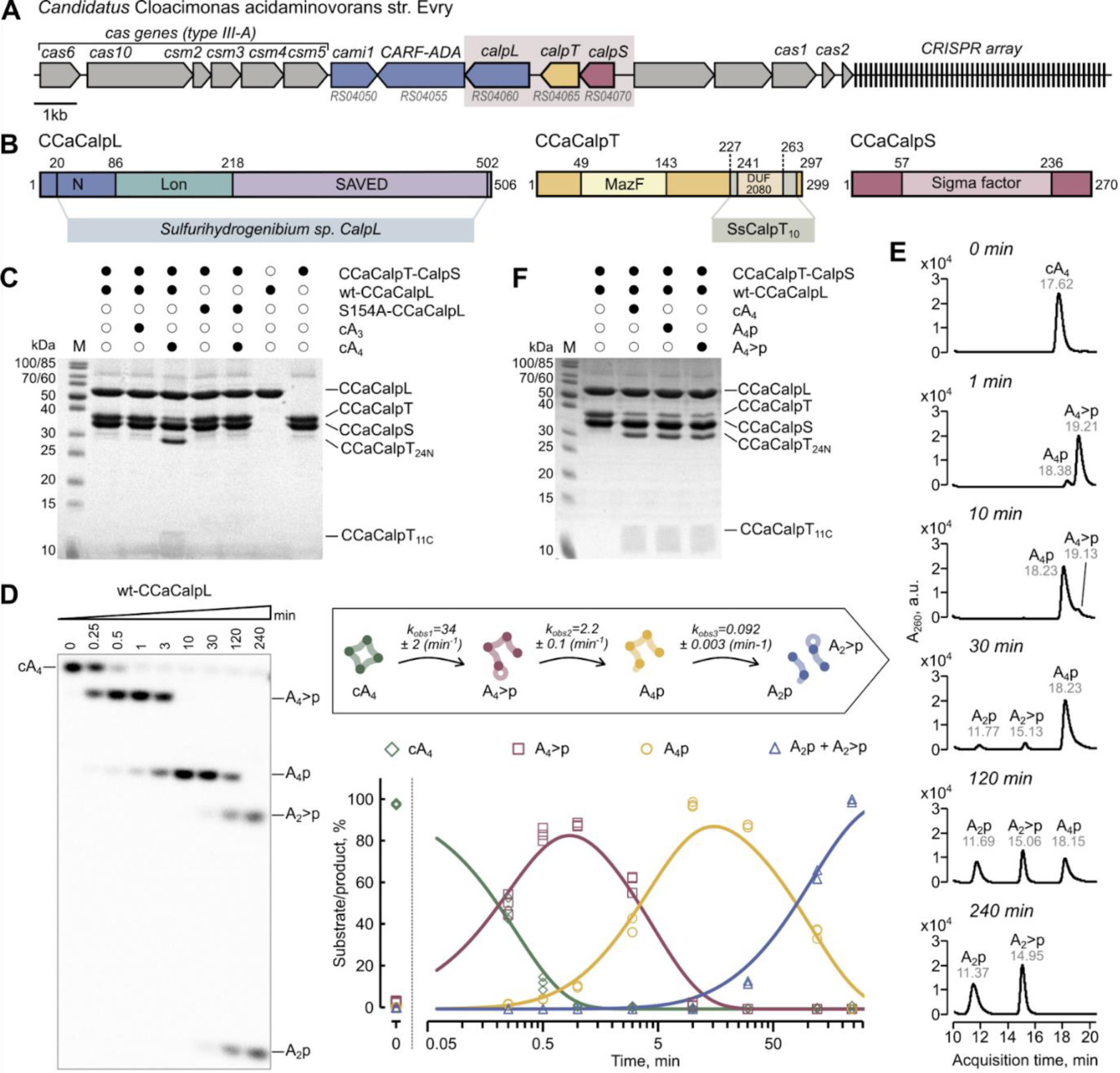
Protease and ring nuclease activities of CCaCalpL. (A) Schematic representation of the *Candidatus* Cloacimonas acidaminovorans (CCa) type III-A CRISPR-Cas locus. Genes of potential associated effector proteins are colored blue. Genes of the tripartite *calpL-calpT-calpS* effector system are shown in the pink background. (B) Schematic representation of CCaCalpL, CCaCalpT, and CCaCalpS with homologous regions identified using HHpred. CCaCalpL shows 46.6% similarity to CalpL from *Sulfurihydrogenibium* spp. YO3AOP1^16^. (C) CCaCalpL protease cleavage reaction analyzed by SDS-PAGE and Coomassie staining. M, protein molecular weight marker. Wt-CCaCalpL and protease active site mutant S154A-CCaCalpL were incubated with CCaCalpT-CalpS heterodimer in the presence of cA_3_, cA_4_, or no cA_n_ for 120 min at 37 °C. The reactions contained 5 µM CCaCalpL, 5 µM CCaCalpT-CalpS heterodimer and 0 or 25 µM cA_4_ or cA_3_. (D) Cleavage of α^32^P-labeled cA_4_ by wt-CCaCalpL under multiple turnover conditions analyzed by denaturing PAGE (left). The reactions contained 1 µM wt-CCaCalpL and 10 µM cA_4_ including 10 nM α^32^P-labeled cA_4_. Time course of substrate cA_4_, intermediate products A_4_>p and A_4_p and final reaction products A_2_>p and A_2_p are plotted (bottom right). Averaged exponential fits (solid lines) to the substrate cleavage data of three experiments are shown. Cartoon depicting the reaction course with indicated observed turnover rates (k_obs1_, k_obs2_ and k_obs3_) is shown above the plot. Turnover rates were calculated as described in Star Methods. (E) Cleavage of cA_4_ by wt-CCaCalpL under multiple turnover conditions analyzed by HPLC-MS. (F) Products of protease reactions performed by wt-CCaCalpL using cA_4_ or intermediate cleavage products A_4_>p and A_4_p as activators. 120 min reactions contained 5 µM CCaCalpL, 5 µM CCaCalpT-CalpS and 0 or 25 µM cA_4_, A_4_>p or A_4_p. Reaction products were analyzed by SDS-PAGE and Coomassie staining. See also Figures S1, S2, S3 and Table S1.

Further, we aimed to elucidate functional and structural mechanisms of the tripartite CCaCalpL-CalpT-CalpS effector system. Since different effectors may be activated by different cA_n_ in the same bacteria^31,32^, we sought to identify the activating cA_n_ by examining cA_n_ production by the native CCaCsm complex *in vitro*. For this purpose, we constructed expression vectors encoding the *E. coli* codon-optimized CCaCsm complex with a minimal CRISPR array (Table S1), expressed the complex in *E. coli,* and purified it by affinity chromatography (Figure S1A). High-performance liquid chromatography-mass spectrometry (HPLC-MS) analysis of cyclase reaction products revealed that CCaCsm produces both cA_3_ and cA_4_, with the major product being cA_4_ (Figure S1B). Next, to obtain the components of the CCaCalpL-CalpT-CalpS effector system, we constructed expression vectors that encode *E. coli* codon-optimized variants of the *CCacalpL* and *CCacalpT-calpS* genes (Table S1). CCaCalp proteins were expressed in *E. coli* and purified using affinity chromatography and size-exclusion chromatography (SEC) (Figure S2A). Subsequent SEC with multi-angle light scattering (SEC-MALS) analysis confirmed that purified CCaCalpL is a monomer in solution and that purified co-expressed CCaCalpT and CCaCalpS form a stable heterodimeric complex (Figure S2B). We further subjected CCaCalpT-CalpS to the CCaCalpL proteolysis reaction in the presence of potential activators cA_3_, cA_4_, or no added cA_n_. SDS-PAGE analysis revealed that cA_4_, but not cA_3_ activated the CCaCalpL protease to cleave the CCaCalpT protein in the CCaCalpT-CalpS heterodimer (Figures 1C, S2C, S2D and S2E). This result is consistent with cA_4_-induced ribosome-dependent mRNA degradation by CCaCami1^11^. Mutation of the conserved S154 to alanine in the predicted Lon protease active site completely abolished the CCaCalpL protease activity (Figure 1C). To identify the cleavage site in CCaCalpT protein, we purified the C-tagged version of a single CCaCalpT protein, subjected it to cleavage by CCaCalpL, and analyzed the cleavage products using HPLC-MS (Figure S2F). We determined that CCaCalpL cleaves CCaCalpT in an alanine rich motive ^202^LAAA between A204 and A205 amino acids resulting in a 24.3 kDa N-terminal fragment (CCaCalpT_24N_) with the MazF domain, and a 10.9 kDa C-terminal fragment (CCaCalpT_11C_) with the DUF2080 domain (Figure S2F). Taken together, these results confirm that CCaCalpL-CalpT-CalpS is a tripartite cA_4_-activated effector system, in which CCaCalpL is an cA_4_-dependent protease that cleaves CCaCalpT in the CCaCalpT-CalpS complex.

### CalpL SAVED domain is a ring nuclease

Recent studies revealed that CARF effectors, which participate in cA_n_-signaling, often possess intrinsic ring nuclease activity^26–29^. The CARF domain of the neighboring effector CCaCami1 also exhibits ring nuclease activity^11^. This led us to hypothesize that type III CRISPR-Cas-associated effectors in *Ca*. C. acidaminovorans are not intended for abortive infection and hence the SAVED domain of CCaCalpL may also degrade its activator cA_4_, in a manner similar to CARF ring nucleases. By incubating CCaCalpL with cA_4_ and analyzing the reaction products by HPLC-MS, we found that CCaCalpL indeed cleaves cA_4_ (Figures S3A). Three cA_4_ cleavage products were detected: A_4_p (ApApApA with a 3’-phosphate), A_2_>p (ApA with a 2′,3′-cyclic phosphate) and A_2_p (ApA with a 3’-phosphate). No differences were observed in cA_4_ cleavage with Mg^2+^, Mn^2+^, or EDTA, indicating that cA_4_ cleavage is a metal-independent reaction (Figure S3B). This demonstrates that CCaCalpL is a metal-independent SAVED ring nuclease specific for cA_4_ degradation.

To elucidate the CCaCalpL ring nuclease mechanism, we analyzed the time course of a multiple turnover ([S] > [E], 10 µM cA_4_ and 1 µM CCaCalpL) cA_4_ cleavage reaction by denaturing PAGE and HPLC-MS (Figures 1D and 1E). We found that radiolabeled cA_4_ is first converted into the first linear reaction intermediate A_4_>p (ApApApA with a 2′,3′-cyclic phosphate), followed by conversion of A_4_>p to the second linear intermediate A_4_p, which is eventually cut into equimolar amounts of linear dinucleotides A_2_>p and A_2_p (Figure 1D).

Accumulation of large amounts (up to 9 µM or up to 90% of all reaction products, i.e. 9-fold higher than CCaCalpL concentration) of both A_4_>p and A_4_p intermediates implies that (i) after each cleavage step the reaction product dissociates from the CCaCalpL enzyme, and (ii) CCaCalpL reaction rates drop by more than 10-fold for each subsequent reaction stage. Indeed, fitting a set of equations describing a three-step consecutive reaction yielded enzyme turnover rate estimates for all three stages: 34 min^-1^ for cA_4_ conversion to A_4_>p, 2.2 min^-1^ for A_4_>p conversion to A_4_p, and 0.10 min^-1^ for A_4_p conversion to A_2_>p + A_2_p (Figure 1D). Under single turnover conditions ([E] > [S], 1 uM CCaCalpL and 50 nM cA_4_) the reaction followed the same mechanism, where cA_4_ was relatively rapidly (on a scale of seconds) converted to A_4_>p and A_4_p, whereas A_4_p was relatively slowly (on a scale of tens of minutes) converted into A_2_>p and A_2_p (Figure S3C).

HPLC-MS analysis confirmed that CCaCalpL can convert both A_4_>p and A_4_p intermediates into the final products A_2_p and A_2_>p (Figure S3D). Furthermore, we found that both A_4_>p and A_4_p stimulate the protease activity of CCaCalpL to a similar extent as cA_4_ (Figure 1F).

### cA_4_-activated CalpL forms filaments

To structurally characterize the CCaCalpL effector, we performed cryogenic electron microscopy (cryo-EM) imaging of CCaCalpL samples in the presence of the activator cA_4_ or its cleavage intermediates A_4_>p or A_4_p (Figure S4 and Table S2). Cryo-EM analysis revealed that CCaCalpL proteins stack together and form ordered short filamentous structures (Figure 2A), resembling several other SAVED domain-containing proteins, including TIR-SAVED and Cap4 from CBASS systems^33,34^, which form filaments upon activator molecule binding.

**Figure 2.**
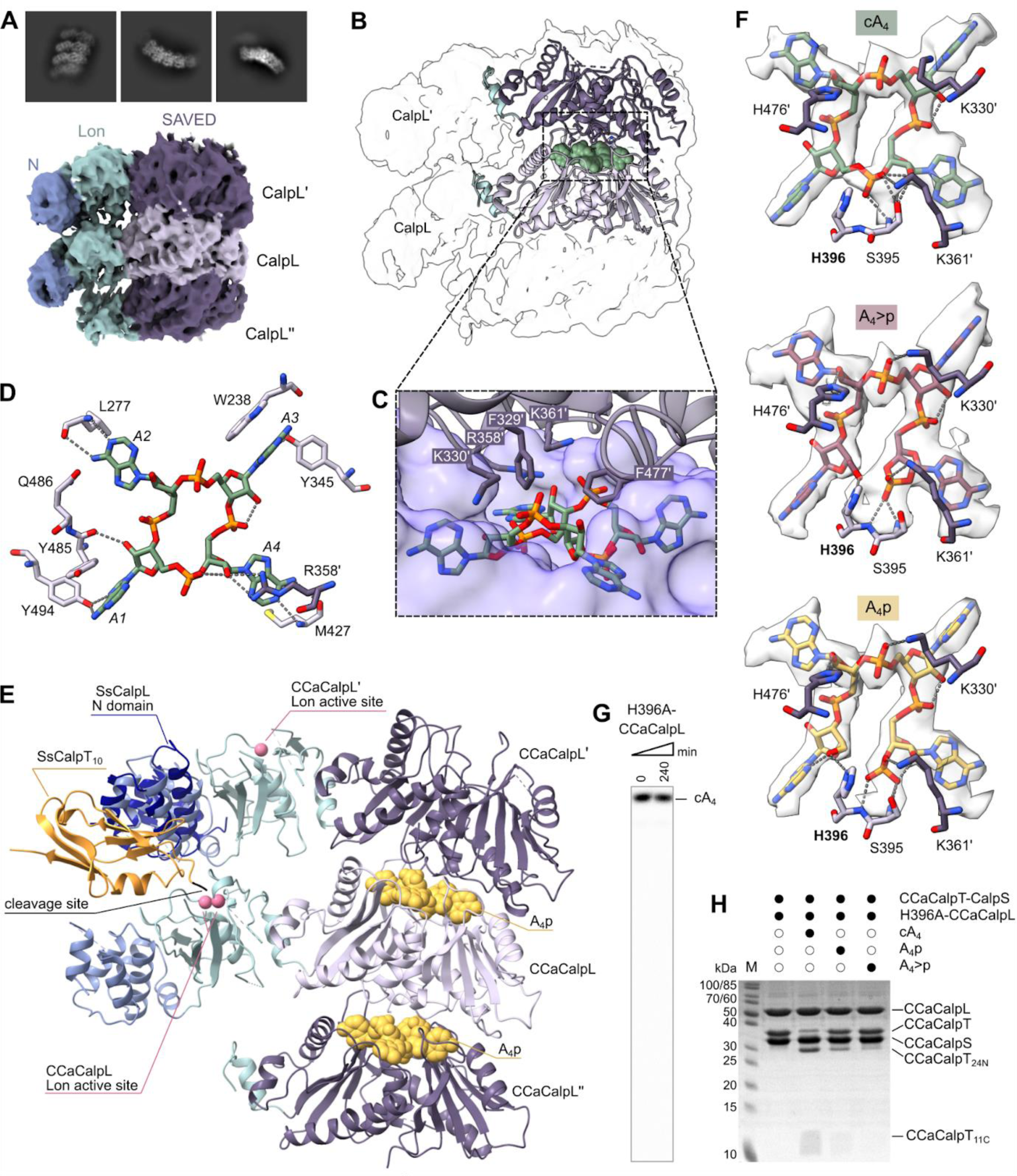
Cryo-EM structure of activated CCaCalpL filaments. (A) Representative 2D classes (top) and electron density map (bottom) of the CCaCalpL filament bound to cA_4_. (B) The atomic model of two SAVED domains (dark purple and light purple) from adjacent CCaCalpL subunits depicted on the electron density map. The cA_4_ (green) is located between SAVED domains. (C) Interactions between neighboring SAVED domains and sandwiched cA_4_. CCaCalpL (light purple) subunit binds cA_4_ in a deep pocket. The adjacent CCaCalpL’ (dark purple) subunit forms fewer contacts to cA_4_. (D) Amino acid residues that form contacts with cA_4_ adenine bases. The residues from the CCaCalpL subunit are depicted in light purple, the residues of the adjacent CCaCalpL’ subunit are shown in dark purple. (E) Superimposition of partial post-cleavage complex SsCalpL N-domain and SsCalpT_10_ (PDB ID: 8B0U) onto A_4_p-bound CCaCalpL filament. The cleavage site of SsCalpT is located in the proximity of Lon active site of the adjacent CCaCalpL monomer. (F) Potentially nucleophilic residues that interact with cA_4_, A_4_>p and A_4_p and may be involved in the ring nuclease activity. (G) Cleavage of α^32^P-labeled cA_4_ by H396A-CCaCalpL under single turnover conditions analyzed by denaturing PAGE. The reactions contained 1 µM wt-CCaCalpL and 50 nM α^32^P-labeled cA_4_. (H) Protease reaction performed with H396A-CCaCalpL using cA_4_ or intermediate cleavage products A_4_>p and A_4_p as activators. 120 min reactions contained 5 µM CCaCalpL, 5 µM CCaCalpT-CalpS and 0 or 25 µM cA_4_, A_4_>p or A_4_p. Reaction products were analyzed by SDS-PAGE and Coomassie staining. See also Figures S4, S5, S6 and Table S2.

The best quality cryo-EM map (2.75 Å resolution) was obtained for the A_4_p-bound CCaCalpL, which allowed us to model two adjacent full-length CCaCalpL subunits and a SAVED domain of the third subunit, with A_4_p molecules bound between the adjacent SAVED domains (Figure S5A). The 2.97 Å map of the cA_4_-bound CCaCalpL and 2.81 Å map of the A_4_>p-bound CCaCalpL allowed us to model only the SAVED domains from two (cA_4_ structure) or three (A_4_>p structure) adjacent CCaCalpL subunits sandwiching the respective activator molecule(s) (Figures 2B and S5A). Superposition of cA_4_/A_4_>p/A_4_p-bound CCaCalpL SAVED domains revealed no significant differences (Figure S5A). In the filament, CCaCalpL monomers oligomerize in a head-to-tail fashion, with the adjacent SAVED domains sandwiching cA_4_/A_4_>p/A_4_p between the opposite surfaces in a manner similar to other SAVED effectors (Figures 2B and 2C). The first (primary) CCaCalpL subunit binds the tetra-adenylate in a deep pocket, while the adjacent (secondary) subunit CCaCalpL’ utilizes a raised patch with two phenylalanine “fingers” on the opposite domain side (Figure 2C). Superposition of bound cA_4_/A_4_>p/A_4_p tetra-adenylates revealed that they are all bound in almost identical conformations (Figure S5B). All four adenine bases are inserted into tight pockets and stabilized by numerous contacts with amino acids from the primary CCaCalpL subunit (Figure 2D). Only R358’ from the adjacent CCaCalpL’ interacts with one of the adenosine bases. The protein also forms extensive contacts with the phosphodiester backbone of the cA_4_/A_4_>p/A_4_p tetra-andenylates (Figure S5C). Taken together, the structures show how sandwiching of cA_4_ or its cleavage products A_4_>p/A_4_p between the SAVED domains enables CCaCalpL oligomerization.

### Mechanism of CCaCalpT cleavage

The overall structure of a single CCaCalpL subunit within the filament is similar to the structure of SsCalpL monomer from a homologous system (PDB ID: 8B0R, DALI Z score 33.1, RMSD=4.1 Å), with the CCaCalpL Lon catalytic residues S154 and K195 coinciding with the SsCalpL catalytic dyad S152-K193 (Figure S5D). In the crystal structure of the SsCalpL-CalpT_10_ post-cleavage complex (PDB ID: 8B0U, SsCalpT_10_ corresponds to CCaCalpT_11C_), the SsCalpT cleavage site is located on the opposite side of SsCalpL from the Lon protease catalytic center, suggesting that binding and cleavage of SsCalpT is performed by two separate SsCalpL proteins, although the exact mechanism of CalpT cleavage by CalpL was not determined^16^. Intriguingly, an overlay of the partial SsCalpL-SsCalpT_10_ post-cleavage complex (SsCalpL N-domain and SsCalpT_10_) and CCaCalpL-A_4_p filament structures places the SsCalpT cleavage site in proximity (approximately 14 Å) from the active site of Lon protease of the adjacent CCaCalpL subunit in the filament (Figure 2E). Such *in trans* CalpT cleavage mechanism, by which cleavage of the bound CalpT is carried out by the Lon protease domain of the adjacent CalpL subunit within the filament, explains the importance of the filament formation for protease activity. Indeed, the observed dependence of CCaCalpL protease activity on cA4 concentration (Figure S2E) suggests that filament formation occurs at micromolar activator concentrations.

### Active site(s) of the SAVED ring nuclease

Based on the structures of cA_4_/A_4_>p/A_4_p-bound CCaCalpL filaments, we have identified residues S395 and H396 as the likely candidates for the ring nuclease catalytic center. Both S395 and H396 are in close proximity to the A1-A4 phosphodiester bond of cA_4_ and the termini of linear A_4_>p and A_4_p intermediates (Figure 2F), suggesting their involvement in the ring opening of cA_4_ (conversion cA_4_ to A_4_>p) and subsequent hydrolysis of the 2′,3′-cyclic phosphate (conversion of A_4_>p to A_4_p).

Mutation S395A significantly reduced the cA_4_ hydrolysis rate (Figure S6A), whereas the H396A replacement completely abolished the cleavage of cA_4_ and linear intermediates A_4_>p and A_4_p, confirming the pivotal role of H396 in catalysis (Figures 2G and S6B). The protease activity of H396A-CCaCalpL was still stimulated by cA_4_, but was severely diminished in the presence of either A_4_>p and A_4_p (Figure 2H). This implies that the mutant retained the ability to bind cA_4_, but not the linear tetra-adenylates, pinpointing the key role of H396 in coordinating the open 5′-OH and 2′,3′-cyclic phosphate/3′-phosphate termini of the A_4_>p and A_4_p intermediates, as observed in the respective CCaCalpL structures (Figure 2F).

The preferred binding orientation of A_4_>p and A_4_p intermediates in respect to the H396 catalytic residue provides a simple explanation for the observed cA_4_ cleavage pattern, which exclusively yields dinucleotides A_2_>p and A_2_p without any A_1_ + A_3_ products (Figures 1D, 1E and S3C). Indeed, if we assume that the CCaCalpL has the second catalytic center at the opposite side of the bound tetra-adenylate (close to the A2-A3 scissile phosphate), ensuing cleavage at this catalytic center would yield the observed dinucleotide products. Relatively slow conversion of A_4_p to A_2_>p + A_2_p (Figure 1D) may be attributed to lower catalytic efficiency of this second catalytic center. The most likely catalytic residues for this reaction are H476’ and K330’ from the adjacent SAVED domain, which are both located in the immediate proximity to the A2-A3 phosphate opposite of H396 (Figure 2F). However, neither of K330 nor H476 mutations abolished formation of the dinucleotide products (Figures S6C, S6D and S6E), ruling out their direct involvement in catalysis. On the other hand, some CARF ring nucleases, such as EiCsm6, have been proposed to degrade cA_6_ primarily through enforcing ligand conformation compatible with an in-line nucleophilic substitution, without direct involvement of any catalytic residues^28^. A similar mechanism may be proposed for the second CCaCalpL SAVED domain active site. This catalytic center, if present, could also cleave cA_4_ in the presence of the H396A mutation that inactivates the first catalytic center. Since this is not the case (Figure 2G), we must assume that the second catalytic center becomes active only when bound to linear A_4_p intermediate, but not the cyclic cA_4_.

Though both the above ‘second active site’-related assumptions, i. e. conformation-dependent catalysis and the ability to discriminate bound cA_4_ from A_4_p, may be attributed to subtle conformational differences between CCaCalpL bound to cA_4_/A_4_p that are not detectable in our structures due to limited resolution (Figure S5B), we must also consider an alternative mechanism involving just a single catalytic site. In this case we must presume that the linear intermediate A_4_p can bind to the SAVED domain in two orientations: (i) with the open 3’-phosphate and 5’-OH termini facing H396 (the preferred orientation, captured in the A_4_p-bound CCaCalpL structure, Figure 2F, bottom), and (ii) the opposite orientation, which would place the uncleaved A2-A3 phosphate in the proximity of H396, and the 3’-phosphate/5’-OH termini in the proximity of K330’ and H476’. Orientation (ii) must be significantly less populated than (i), accounting for the low cleavage rate of the A_4_p intermediate at the A2-A3 phosphate (Figure 1D), and lack of density corresponding to an uncleaved A2-A3 phosphodiester bond in the proximity of H396 in the A_4_p-bound CCaCalpL structure (Figure 2F, bottom). Nevertheless, the dynamic nature of the CCaCalpL filaments, which allows dissociation of cA_4_ cleavage intermediates from the enzyme during the ring nuclease reaction (as discussed above, Figure 1D), may allow multiple rounds of A_4_p dissociation and re-binding required for occasional A_4_p binding in the (ii) orientation leading to its cleavage. In summary, we demonstrate here that CCaCalpL H396 is the key catalytic residue responsible for the cA_4_ ring opening reaction; the same catalytic center may also be responsible for A_4_p cleavage to the final products A_2_>p and A_2_p, albeit we cannot completely rule out the presence of a second catalytic center.

### Both protease and ring nuclease activities rely on CCaCalpL filament formation

To test the importance of filament formation for CalpL activities, we attempted to disrupt the SAVED domain inter-subunit interface by introducing two charge reversal mutations R358E/K361E. As expected, cryo-EM analysis of the R358E/K361E-CCaCalpL-cA_4_ sample did not reveal any filament particles. R358E/K361E mutations also abolished CCaCalpT cleavage (Figure S6F), confirming that filament formation is indeed required for the protease activity. Next, we used biolayer interferometry (BLI) to test if CCaCalpL filament is required for the CCaCalpT-CalpS target binding (Figure 3A). CCaCalpT-CalpS heterodimer was immobilized on a Ni^2+^-nitrilotriacetic acid (Ni^2+^-NTA) sensor chip as the ligand, while two CCaCalpL variants, the ring nuclease-deficient H396A and filament-deficient R358E/K361E, were used as analytes. No CCaCalpL binding to the sensor surface with immobilized CCaCalpT-CalpS was detected in the absence of cA_4_ with either mutant, but addition of cA_4_ stimulated binding of H396A-CCaCalpL, which was manifested as a significant increase in the BLI signal (Figures 3A and S6G). By contrast, no binding was detected with the R358E/K361E mutant, providing further evidence that CCaCalpL filament formation is essential for its interaction with CCaCalpT-CalpS (Figure 3A).

**Figure 3.**
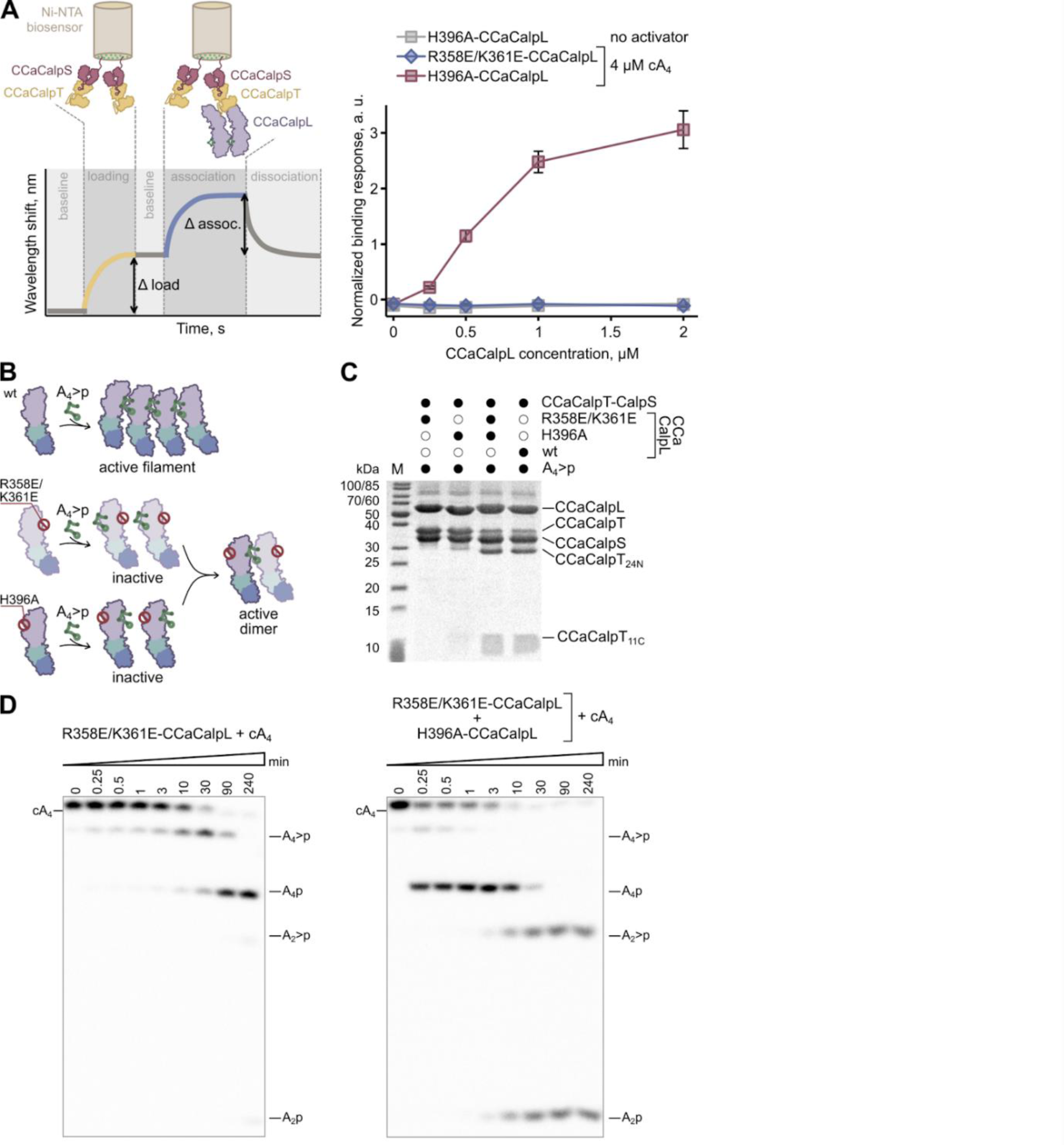
Oligomerization is required both for CCaCalpL protease and ring nuclease activities. (A) Setup of the BLI experiment (left). CCaCalpT-CalpS heterodimer was immobilized onto a Ni-NTA biosensor (loading step). The loaded sensor was used to monitor CCaCalpL binding at various concentrations (association step). Binding experiment results (right). To avoid rapid cA_4_ cleavage during the assay H396A ring nuclease mutant was used instead of wt-CCaCalpL. Normalized binding response equals (Δ association) / (Δ loading). All data points are presented as mean values from three experiments ± 1 SD. (B) Schematic representation of the assembly of the A_4_>p-activated CCaCalpL dimer. (C) Protease reactions were analyzed by SDS-PAGE of the CCaCalpL variants shown in (B). The reactions contained 5 µM CCaCalpL, 5 µM CCaCalpT-CalpS and 25 µM A_4_>p. (D) α^32^P-labeled cA_4_ cleavage reactions by R358/K361E-CCaCalpL or by R358/K361E-CCaCalpL mixed with H396A-CCaCalpL under single turnover conditions. Samples were analyzed by denaturing PAGE. The reaction contained 1 µM CCaCalpL and 50 nM cA_4_. See also Figure S6.

To test if CCaCalpL dimer, i. e. the minimal fragment of a filament, is sufficient for the Lon protease activity, we attempted to reconstitute an active CCaCalpL heterodimer from two inactive variants carrying mutations at the opposite surfaces of the SAVED domain (Figure 3B): (i) the H396A mutant, which is unable to bind A_4_>p and cleave CCaCalpT (Figures 2H and S6B), and (ii) the filament deficient R358E/K361E mutant. Although each individual mutant lacked protease activity, their equimolar mix yielded an active CCaCalpL variant with wt-like CCaCalpT-CalpS cleavage activity (Figure 3C). This finding is also consistent with the *in trans* CCaCalpL cleavage mechanism discussed above (Figure 2E).

The R358E/K361E interface mutations also significantly reduced ring nuclease activity of CCaCalpL (>100-fold compared to wt-CCaCalpL), confirming that filament formation is also required for the ring nuclease activity (Figure 3D, left). As with the protease activity, combining the ring nuclease-deficient mutant H396A with the low activity mutant R358E/K361E restored cA_4_ cleavage activity to a wt-like level (Figures 3D, right, and S3C). The small amount of uncleaved substrate observed over the first minutes of the reaction is likely due to cA_4_ sequestration by the inactive H396A homo-oligomers. Collectively, our findings demonstrate that CCaCalpL oligomerization is crucial for its interaction with the CCaCalpT-CalpS target, and both its protease and ring nuclease activities.

### After cleavage CalpT requires further processing to release CalpS

Recently, it has been demonstrated that CalpS functions as a σ factor by forming a complex with DNA-directed RNA polymerase (RNAP)^16^. To analyze the possible interaction of CCaCalpS with RNAP, first we performed AlphaFold modeling of CCaCalpS and superimposed it onto σ^E^ in the *E. coli* RNAP/σ^E^ transcription initiation complex (PDB: 6JBQ)^35^. The N- and C-terminal domains of CCaCalpS aligned with the promoter binding domains σ_2_ and σ_4_ of σ^E^, respectively (Figure S7A). Then to assess CCaCalpS interaction with RNAP, we performed pull-down experiments from *E. coli* overexpressing the C-terminally tagged CCaCalpS protein. We found that neither expression of CalpS alone, nor CalpS in the CalpT-CalpS complex resulted in co-purification of *E. coli* RNAP (EcRNAP) with CalpS (Figure S2A).

Subsequently, we investigated the CCaCalpS-EcRNAP interaction using *in vitro* assays. Initially, we incubated the purified EcRNAP core enzyme with the C-tagged CCaCalpS to form the RNAP holoenzyme. Magnetic beads were employed to pull down the formed complexes and SDS PAGE analysis of the elution fractions revealed co-purification of the β, β’ and α subunits of EcRNAP with CCaCalpS (Figure S7B). To obtain a more robust binding signal, we utilized the BLI technique (Figure 4A). We immobilized CCaCalpS on the Ni^2+^-NTA biosensor and exposed it to the EcRNAP core enzyme, which resulted in a wavelength shift, confirming the interaction between CalpS and EcRNAP. No EcRNAP binding occurred when the full-length CCaCalpT-CalpS heterodimer or pre-cleaved CCaCalpT-CalpS heterodimer was immobilized on the biosensor. The AlphaFold model of the CCaCalpT-CalpS heterodimer suggests that CCaCalpT interacts with CCaCalpS exclusively with the CCaCalpT_24N_, and the CalpL cleavage site in CalpT is distant from the CCaCalpT-CalpS interaction interface (Figure S2C). We hypothesized that cleavage alone may not be sufficient to disrupt the interaction of the proteins in the complex. Similarly, a stable intact SsCalpT_23_-CalpS complex was observed after the SsCalpL reaction in the homologous system^16^. Therefore, we further tested whether CCaCalpS is released from the heterodimer after the CCaCalpT cleavage using magnetic beads to pull down the proteins that remain associated with CCaCalpS (Figure S8A). Remarkably, CCaCalpS remained bound to both CCaCalpT_24N_ and CCaCalpT_11C_, confirming that the CCaCalpT probably requires further processing in the cell to release CCaCalpS, which can then bind to RNAP.

**Figure 4.**
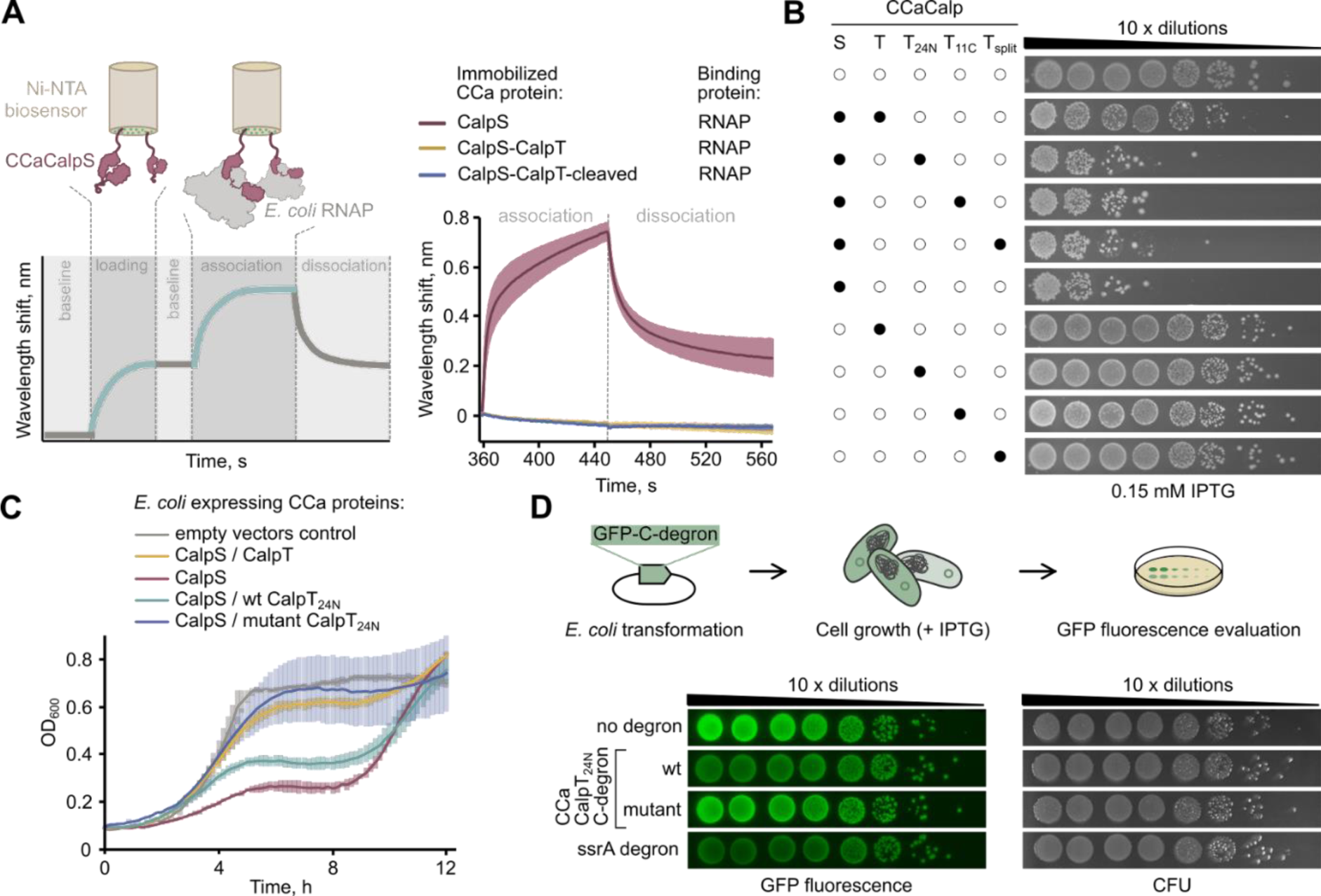
Toxicity of CCaCalpS in *E. coli*. (A) BLI experimental scheme of CCaCalpS binding to *E. coli* RNAP (left). CCaCalpS, CCaCalpT-CalpS heterodimer or CCaCalpT-CalpS heterodimer after cleavage with CCaCalpL were immobilized onto Ni-NTA biosensor (loading step). The biosensor was then used to monitor *E. coli* RNAP binding (association step). Curves of association and dissociation steps of the BLI experiment are shown (right). All data points are presented as mean values from three experiments ± 1 SD. (B) Serial dilutions of *E. coli* expressing combinations of CCaCalpS with full-length CCaCalpT, CCaCalpT_24N_, CCaCalpT_11C_ or split CCaCalpT plated on LB media with IPTG. See Figure S8B for the growth results on LB medium with glucose. (C) Growth curves of *E. coli* expressing CCaCalpS and wt-CCaCalpT_24N_ or mutant CCaCalpT_24N_ with A204D mutation in the predicted C-degron PETLQLAA^204^. (D) GFP fluorescence interference assay in *E. coli*. Experimental workflow (top): *E. coli* cells were transformed with plasmids encoding GFP fused with different degrons: PETLQLAA (CCaCalpT_24N_ C-degron), PETLQLAD (mutant version of CCaCalpT_24N_ C-degron) and ENYALAA (partial fragment of ssrA C-degron). Overnight cultures were serially diluted (10×) and plated on LB medium with IPTG. CCaCalpT_24N_ C-degron resulted in reduced GFP fluorescence (bottom left) without affecting CFU (bottom right). See also Figures S2, S7, S8 and Table S1.

### C-degron of CalpT_24N_ targets it for degradation by cellular proteases

To explore the impact of CCaCalpL-mediated cleavage of CCaCalpT within the CCaCalpT-CalpS system and gain insights into potential CalpT processing in bacterial cells, we performed a *E. coli* survivability assay. For this, we constructed IPTG-inducible vector system encoding *CCacalpS* and a copy of either (i) the full-length *CCacalpT*, (ii) the *CCacalpT_24N_*, (iii) the *CCacalpT_11C_*, or (iv) a split version of *CCacalpT*, which would be formed after its cleavage with CCaCalpL (Table S1). We co-transformed *E. coli* with pairs of vectors, performed serial dilutions and plated cells on LB media with either IPTG or glucose. We observed that *E. coli* expressing CCaCalpS alone formed fewer colonies, while co-expression of full-length CCaCalpT reduced CCaCalpS toxicity (Figures 4B and S8B). No toxicity was observed when CCaCalpT or its cleavage variants were expressed alone, indicating that CCaCalpS is a toxic component of the CCaCalpT-CalpS pair. This aligns with the role of CCaCalpT as an anti-σ factor, preventing the interaction between CCaCalpS σ factor and RNAP.

Interestingly, the cleavage products of CCaCalpT had no effect on CCaCalpS toxicity (Figures 4B). This implies that, in contrast to the *in vitro* experiments, CCaCalpS is released from the CCaCalpT cleavage products *in vivo*. Consequently, we hypothesized that the CCaCalpT_24N_ may carry a degron at its newly formed C-terminus, which targets it for degradation by cellular proteases^36^, in turn releasing CCaCalpS from the heterodimer complex. Based on the AlphaFold model (Figure S2C), we predicted that the terminal 9 amino acids ^196^PEDTLQLAA^204^ (Figure S2F) of CCaCalpT_24N_ are poorly structured and may comprise the C-degron, as unstructured peptides ending with alanines at the C-terminus are commonly found in degrons that target proteins for degradation by the ClpXP unfoldase-protease complex^37–39^.

In bacteria, stalled ribosome rescue systems add a C-terminal poly-alanine degron^40^ or a C-terminal SsrA-degron ending in LAA to proteins stuck in stalled ribosomes, directing them for degradation by ClpXP^17,41^. The terminal AA motif is crucial for C-degron recognition, and mutating either alanine to aspartate in the terminal AA motif significantly reduces efficiency of ClpXP degradation^42^. Therefore, to test our CCaCalpT_24N_ C-degron hypothesis, we mutated the C-terminal A204 residue in the putative CCaCalpT_24N_ C-degron to aspartate and examined its impact on the CCaCalpS toxicity in *E. coli* by registering growth curves in liquid LB after IPTG induction (Figures 4C). Cells expressing CCaCalpS alone or co-expressing CCaCalpS with the wt CCaCalpT_24N_ exhibited poor growth. However, co-expression of CCaCalpS with the CCaCalpT_24N_ with mutated degron resulted in normal *E. coli* growth, comparable to cells co-expressing CCaCalpS and full-length CCaCalpT (Figure 4C). We also employed the GFP reporter system, using expression vectors encoding *gfp* fused at the C-terminus with: (i) the wt C-degron of CCaCalpT_24N_ (PEDTLQLAA), (ii) the mutant C-degron of CCaCalpT_24N_ (PEDTLQLAD), and (iii) a partial SsrA-C-degron (ENYALAA)^37^ as a positive control (Table S1). *E. coli* cells were transformed with these vectors and serial dilutions of cells were plated onto LB media with IPTG (Figure 4D). While cell survivability was not affected, a decrease in GFP fluorescence was observed in cells expressing GFP fused with wt C-degron, but not with mutant C-degron. Furthermore, we confirmed the results of our spot assay by quantitatively analyzing the fluorescence levels of GFP-C-degron in liquid culture over time (Figure S8C). Taken together, these results demonstrate that CCaCalpL cleavage product CCaCalpT_24N_ undergoes further degradation by cellular proteases. Only after the degradation of CCaCalpT_24N_, CCaCalpS is released from the heterodimer, causing growth arrest in *E. coli*.

## DISCUSSION

### Mechanism of CalpL-CalpT-CalpS cascade

In this study, we demonstrate that the tripartite CalpL-CalpT-CalpS effector system releases extracytoplasmic function (ECF)-like σ factor, CalpS, in response to the cA_4_ signal produced by Csm complex associated with the type III CRISPR-Cas system. The proposed model for the type III CRISPR-Cas CalpL-CalpT-CalpS-mediated immunity is summarized in Figure 5. Initially, cA_4_ produced by the type III Csm/Cmr complex binds to the SAVED domain of the CalpL monomer, inducing formation of a CalpL filament and activating the CalpL protease. This filament, in turn, recruits the CalpT-CalpS anti-σ factor/σ heterodimer, leading to cleavage of CalpT. Within the CalpL filament, one CalpL subunit binds CalpT and the adjacent CalpL subunit cleaves it. Following hydrolysis, both cleavage products CalpT_24N_ and CalpT_11C_ remain bound in the complex with CalpS. However, cleavage exposes the C-degron of CalpT_24N_, which directs it for degradation by cellular proteases. This, in turn, releases the σ factor CalpS, which binds to RNAP, thereby altering transcription and allowing the cell to survive infection. The CalpL serves as a self-limiting effector, with its SAVED domain acting as a ring nuclease that cleaves its activator cA_4_. The relatively rapidly formed intermediates A_4_>p and A_4_p both stimulate protease activity and preserve filament structure of CalpL. Such sequential cleavage of cA_4_ activator by CCaCalpL thus serves as a “timer” that limits the duration of the activated state of the effector, similar to Csm6 and Cami1^11,26,29^.

**Figure 5.**
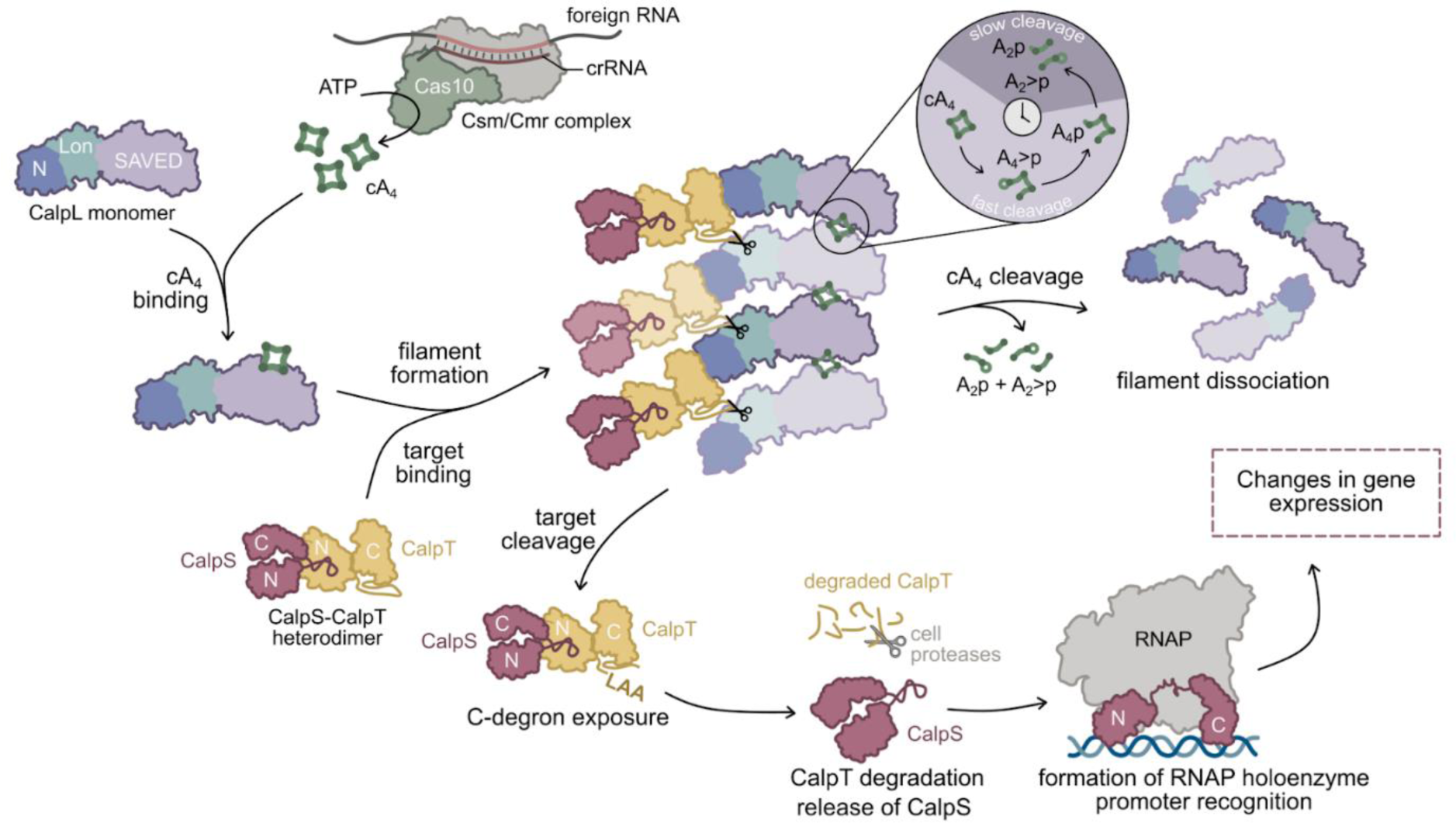
Mechanism of the antiviral defense mediated by the cA_4_-activated tripartite CalpL-CalpT-CalpS system. Binding of foreign RNA triggers cA_4_ synthesis by the type III CRISPR-Cas complex. Effector CalpL uses a sensory domain SAVED to bind cA_4_. cA_4_ binding induces filament formation, which enables cleavage of CalpT in the CalpL-bound CalpT-CalpS heterodimer by the CalpL Lon domain from an adjacent CalpL subunit. CalpT cleavage leads to the subsequent CalpT degradation by cellular proteases liberating CalpS to bind to RNA polymerase and act as a σ factor for the transcription of specific genes. The ring nuclease activity of the SAVED domain regulates the Lon protease activity of CalpL by cleaving its activator cA_4_.

Temporal and conditional control of transcription initiation is a common mechanism by which bacteria counteract stress or adapt to changing ecological conditions. Many bacteria possess alternative σ-factors that redirect RNAP to genes necessary for adaptive responses. Often, alternative σ factors are controlled by specific anti-σ proteins^43^. The σ factor is typically released in response to a signal perceived by either the anti-σ factor or other components in more complex signal relay systems. Here we showed that cleavage of the anti-σ factor CalpT by the CalpL protease alone is not sufficient to release the ECF-like σ factor CalpS from the CalpT-CalpS heterodimer. We found that cA_4_-activated CalpL Lon protease recognizes and cleaves a poorly structured region of CalpT and the newly formed C-terminus of CalpT_24N_ exposes LAA-ending degron. This directs CalpT_24N_ for further degradation by cellular proteases in *E. coli*. This resembles other ECF-like sigma factors, e.g. RseA-σ^E^ pair, where the anti-σ RseA is cleaved to expose C-terminal degron (VAA), which targets it for degradation by the cytoplasmic AAA+ protease ClpXP, liberating σ^E^ ^44–46^.

### CalpL SAVED domain is a ring nuclease

Ring nucleases characterized to date are symmetric homodimers that utilize two identical active sites to split cA_4_ or cA_6_ rings into two halves^23^. For example, the dimeric CARF-fold Crn1 catalyze in-line nucleophilic attacks of two opposite cA_4_ 2′-OH groups on the adjacent phosphodiester bonds, yielding A_2_>p products with 2’,3’-cyclic phosphates. The reactions on the opposite sides of the cA_4_ ring are not concerted, but the linearized intermediate A_4_>p is not released from the active site^47–49^. A similar mechanism is employed by CARF domains of the self-limiting CARF effectors that cut cA_4_ or cA_6_ to A_2_>p and A_3_>p, respectively^26–29^. CARF-unrelated Crn2 ring nucleases also cleave cA_4_ into two A_2_>p molecules, but they additionally hydrolyze the cyclic phosphates yielding A_2_p as the final products^50^. We show here that CCaCalpL ring nuclease combines mechanistic features of both Crn1 and Crn2, as it catalyzes both the attack of 2’-OH on the adjacent phosphodiester bond (conversions cA_4_ to A_4_>p and A_4_p to A_2_>p + A_2_p), and hydrolysis of the cyclic phosphate (conversion A_4_>p to A_4_p). The most notable differences of CCaCalpL ring nuclease from Crn1/Crn2 is (i) the asymmetry of its cA_4_ binding cleft, which is formed between two SAVED domains arranged in a head-to-tail fashion, and (ii) sequential manner of cA_4_ cleavage with dissociation of linear A_4_ intermediates from the enzyme. In order to convert cA_4_ into dinucleotide products, an asymmetric CCaCalpL SAVED dimer must either employ two non-identical catalytic centers on the opposite cA_4_ phosphates, or cut these phosphates via repeated use of a single catalytic center. The latter mechanism must involve dissociation of the A_4_p intermediate and its re-binding to the enzyme in the opposite orientation, thereby explaining the relatively rapid release of reaction intermediates from the CCaCalpL active site into solution (Figure 1D).

Notably, similarly to CCaCalpL, the STAS (Sulfate Transporter and Anti-Sigma factor antagonist) fold Crn3 family ring nucleases also form filaments upon cA_4_ binding, which is obligatory for ring nuclease activity^25^, however symmetric active site of Crn3 cleaves both cA_4_ and polyA RNA and requires metal ions for its activity^51,52^, whereas SAVED is a metal-independent cA_4_ ring nuclease.

### Filament formation as a general mechanism for prokaryotic immune effectors

The head-to-tail association of SAVED domains via a signaling molecule observed in the CCaCalpL filament appears to be a conserved feature of all SAVED domain-containing prokaryotic effectors. SAVED filament formation may lead to the assembly of effector’s composite active site, as demonstrated for the CBASS-associated TIR-SAVED^33^, or the effector domain may be allosterically activated via interdomain interactions within the cA_3_-mediated SAVED filament, as described for the CRISPR type III-associated SAVED-CHAT protein^53^. Formation of short filaments of the DUF4297-SAVED fusion protein Cap4 brings together nuclease-like DUF4297 domains, thereby enabling cleavage of double stranded DNA^34^. Distinctly, the HNH-SAVED fusion protein Cap5 in the absence of a signaling molecule forms symmetric HNH-domain swapped dimers with limited contacts between the SAVED domains, but activator binding leads to head-to-tail stacking of SAVED domains within the dimer, or even further oligomerization into tetramers, positioning two HNH sites for double stranded DNA cleavage^54,55^. Our structural analysis of the cA_4_-mediated CCaCalpL filament suggests yet another activation mechanism, by which filament formation makes the CalpT-CalpS substrate bound to one CCaCalpL subunit accessible for the protease active site of the adjacent subunit within the filament.

Activation of effector proteins through filament formation is widespread in prokaryotic antiviral defense systems, as exemplified by TIR-STING effector of CBASS, BcPycTIR effector of Pycsar, and SIR2-SLOG effector of the Thoeris system^56–58^. They usually form long highly stable activated filaments, as it was demonstrated for the SIR2-SLOG effector of the Thoeris system^58^ and are therefore attributed to abortive infection systems^59^. In contrast, our data suggests that CCaCalpL filaments are highly dynamic, as the adjacent SAVED domains bound to cA_4_ must disassemble after each ring nuclease cleavage step, enabling release of the intermediate products. This may increase the chance of recovery for the infected cell after the infection has been defeated and the signaling molecules are cleaved by CCaCalpL.

## MATERIALS AND METHODS

### Homology search

The HHPred tool^60^ was used to search for homology of *Ca.* C. acidaminovorans proteins of interest (NCBI Reference Sequences: WP_044278915.1, WP_015424588.1, WP_015424587.1, WP_015424586.1) in structural and domain databases: PDB, Pfam-A. COG_KOG, NCBI (CD). The search was performed with default settings.

The DaliSearch tool^61^ was used to search for remote structural homologs of the CCaCalpL SAVED domain and full-length CCaCalpL. The SAVED domain (219-503 aa) from the solved cA_4_-bound CCaCalpL filament structure and a single CCaCalpL subunit from the A4p-bound CCaCalpL filament structure were used as a queries.

### Structure prediction

The CCaCalpS protein and the CCaCalpT-CalpS heterodimer were modeled with AlphaFold^62,63^ under the ColabFold^64^ framework using default parameters.

### Cloning and mutagenesis

The coding sequences of the *Candidatus* Cloacimonas acidaminovorans strain Evry proteins of interest and the repeat sequence, were obtained from the NCBI reference genome sequence NC_020449.1. All gene sequences of the proteins have been optimized for expression in *E. coli*. The *E. coli* DH5α strain was used for cloning and plasmid amplification. All vectors constructed in this study are listed in Table S1.

Synthetic gene fragments encoding *cas10* (fused to TEV-cleavable His_6_-StrepII-His_6_ N-tag), *csm2*, *csm3*, *csm4* and *csm5* genes were purchased from Twist Bioscience (United States) and cloned into the pCDFDuet-1 plasmid via Gibson assembly to obtain pCsm plasmid. The pCRISPR plasmid encoding the minimal CRISPR region (repeat-spacer-repeat) and *cas6* gene was constructed by cloning a synthetic gene fragment into pETDuet-1 backbone using Gibson assembly. The gene fragment encoding the minimal CRISPR region was purchased from GenScript Biotech (United States), and the gene fragment encoding *cas6* was purchased from Twist Bioscience. The DNA synthesis of *calpL* fused with TEV-cleavable His_6_-StrepII-His_6_ tag at the N- or C-terminal and cloning into pBAD/HisA vector (Invitrogen) was ordered from Twist Bioscience. To generate plasmids encoding mutant versions of c*alpL*, site-directed mutagenesis was used with primers carrying the desired mutations. Synthetic gene fragments encoding *calpS* and *calpT* genes were purchased from Twist Bioscience and were used as PCR templates to generate desired fragments for further cloning. The Gibson assembly strategy was used to clone *calpS* into pETDuet-1 vector, which fuses the protein with TEV-cleavable His_6_-StrepII-His_6_ tag at the C-terminus (pET-CalpS-Tag). Similarly, the pET-CalpT-Tag, a pETDuet-1 vector encoding *calpT* fused to a His_6_-StrepII-His_6_ tag at the C-terminus, was generated. Gibson assembly was also used to generate pCDF-CalpT, a pCDFDuet-1 vector encoding *calpT* without the tag.

For the *E. coli* survivability assay pCalpS, pCalpT, pCalpT_24N_, pCalpT_11C_ and pCalpT_split_ vectors were generated. The pCalpS vector was produced by cloning the *calpS* gene into the pCDFDuet-1 vector using the Gibson assembly method. The pCalpT vector encodes the entire *calpT* gene spanning 1-299 codons and was generated by site-directed mutagenesis of the pET-CalpT-tag vector, excluding the segment encoding the affinity tag. The pCalpT_24N_ vector encodes the first 204 codons of the *calpT* gene, while the pCCaCalpT_11C_ vector encodes codons 205-299 (with an additional methionine for initiation) of the *calpT* gene. These vectors were generated from the pCalpT plasmid using site-directed mutagenesis to remove the corresponding parts of the *calpT* gene. The pCalpT_split_ plasmid encodes codons 1-204 (CalpT_24N_) and 205-299 (CalpT_11C_) of the *calpT* gene, separated by a 45 nt spacer encoding a ribosome binding site and an additional methionine as an initiation codon for the expression of the CalpT_11C_. To produce the pCalpT_split_ plasmid, a whole plasmid PCR fragment was generated from the pCalpT template and assembled with a 77 nt bridging oligo using the Gibson assembly method.

The pCalpT_24N_(mut) plasmid encodes the first 204 codons of CalpT with an A204D mutation at the end of the potential degron sequence and was constructed by site-directed mutagenesis using pCalpT_24N_ as a template. pGFP was constructed by cloning the *GFPmut1*^65^ gene into pETDuet-1 vector via the Gibson assembly. Subsequently, pGFP was used as a template in a site-directed mutagenesis strategy to generate plasmids encoding GFP fused to degron sequences: (i) wt CalpT_24N_ C-degron PETLQLAA, (ii) mutant CalpT_24N_ C-degron PETLQLAD, or (iii) partial ssrA C-degron ENYALAA, generating pGFP-(wt)CalpT_24N_-degron, pGFP-(mut)CalpT_24N_-degron and pGFP-ssrA-degron plasmids, respectively. Sequence integrity of all constructed plasmids was confirmed by either Sanger sequencing or whole plasmid sequencing (SeqVision, Lithuania).

### Expression and purification of proteins

#### CCaCsm complex purification

The *E. coli* BL21(DE3) strain was co-transformed with pCsm and pCRISPR plasmids. Bacteria were grown at 37°C in the liquid LB medium supplemented with streptomycin (25 µg/ml) and ampicillin (50 µg/ml) until mid-log phase. Expression was induced with 0.5 mM IPTG and the cell suspension was further cultured at 16°C overnight. Cells were harvested and disrupted by sonication, soluble proteins were captured on the StrepTrapXT column (Cytiva) using chromatography buffer A1 (20 mM Tris-HCl (pH 7.5), 0.5 M NaCl, 8 mM 2-mercaptoethanol) and 50 mM biotin in chromatography buffer A1 for elution. To further remove the affinity tag from the Cas10 subunit, the eluted fractions were pooled and incubated with TEV protease^66^ (at a ratio ∼1:25 (w/w) protease:target protein) and 2 mM DTT at 4°C overnight. After the TEV cleavage reaction, the protein sample was buffer exchanged to low salt buffer (20 mM Tris-HCl (pH 7.5), 50 mM NaCl, 8 mM 2-mercaptoethanol) and further purified on a HiTrap Heparin HP column (Cytiva). The target complex was eluted using a 50–1000 mM NaCl gradient. Peak fractions of the eluted sample were dialyzed against storage buffer 10 mM Tris-HCl (pH 7.5), 300 mM NaCl, 1 mM DTT, 0.1 mM EDTA and 50% (v/v) glycerol, and stored at −20°C. The protein composition of the isolated CCaCsm complex was analyzed by SDS-PAGE. The crRNA of the purified CCaCsm complex was extracted and visualized as described for the homologous Csm complex^11^.

#### CCaCalpL purification

Wt and mutant variants of C-tagged CCaCalpL were expressed in the *E. coli* strain ER2566 using the appropriate expression vector. Bacteria were grown at 37°C in LB medium supplemented with ampicillin (100 µg/ml) to the mid-log phase. Expression was induced with 0.2% arabinose and cells were further cultured at 37°C for 4 hours. Cells were harvested by centrifugation and disrupted by sonication. Soluble proteins were captured on the HisTrap HP column (Cytiva) using chromatography buffer A2 (20 mM Tris-HCl (pH 8.0), 0.5 M NaCl, 8 mM 2-mercaptoethanol) and a 5-500 mM imidazole gradient. The eluted fractions were pooled and further purified on the StrepTrapXT column (Cytiva). The target protein was eluted using 50 mM biotin in A2 chromatography buffer. The eluted fractions were pooled, and the affinity tag was removed as described above for the CCaCsm complex. The cleaved tag and TEV protease were removed by His affinity chromatography. Flow-through proteins were dialyzed against a storage buffer 10 mM Tris-HCl (pH 8.0), 200 mM NaCl, 1 mM DTT, 0.1 mM EDTA and 50% (v/v) glycerol and stored at -20°C. The N-tagged CCaCalpL variant was expressed in *E. coli* DH10B strain and purified as described for C-tagged CCaCalpL, except for the TEV cleavage step.

#### Purification of CCaCalpT-CalpS heterodimer

For the production of the CCaCalpT-CalpS heterodimer, the *E. coli* strain ER2566 was co-transformed with the plasmids pET-CalpS-Tag and pCDF-CalpT. Bacteria were grown at 37°C in LB medium supplemented with streptomycin (25 µg/ml) and ampicillin (50 µg/ml) to the mid-log phase. Expression was induced with 0.5 mM IPTG and the cell suspension was further cultured at 37°C for 4 hours. Cells were harvested and disrupted by sonication. Soluble proteins were captured on the HisTrapHP column (Cytiva) using chromatography buffer A1 and a 5-500 mM imidazole gradient. Eluted fractions were pooled and further purified using the StrepTrapXT column (Cytiva) and chromatography buffer A1. The target protein was eluted using 50 mM biotin in A1 chromatography buffer. The eluted fractions were pooled and further purified by size exclusion chromatography using a Superdex 200 10/300 GL column (Cytiva) and chromatography buffer A1. Peak fractions of the eluted sample were dialyzed against a 25 mM Tris-HCl (pH 7.5) buffer containing 250 mM NaCl, 1 mM DTT and 50% (v/v) glycerol, and stored at -20°C.

#### Purification of CCaCalpT and CCaCalpS

CCaCalpT was purified as described for the CCaCalpT-CalpS dimer, except that *E. coli* strain ER2566 was transformed with a single pET-CalpT-Tag plasmid. CCaCalpS was purified as described for the CCaCalpT-CalpS dimer, except that *E. coli* strain ER2566 was transformed with a single pET-CalpS-Tag plasmid and the size exclusion chromatography step was omitted due to the low yield.

### SEC-MALS analysis

SEC-MALS analysis was performed on an HPLC system (Waters Breeze) combined with a miniDAWN MALS detector (Wyatt) and an Optilab refractive index detector (Wyatt). CCaCalpL, CCaCalpT-CalpS, CCaCalpT or CCaCalpS protein samples were diluted with SEC-MALS buffer (20 Tris-HCl (pH 8.0), 100 NaCl, 1 mM DTT) to a final concentration of 0.3 mg/mL. 200 µL of diluted samples were loaded to a Superdex 200 Increase 10/300 GL column (Cytiva) equilibrated with SEC-MALS buffer. Data acquisition and evaluation were performed using ASTRA 7 software (Wyatt).

### Protease assay

A protease reaction was performed by pre-incubating 5 µM of CCaCalpL (when CCaCalpL mutant was combined in the oligomerization assay, 2.5 uM of each mutant was used) and 5 uM of CCaCalpT-CalpS dimer (or CCaCalpT monomer) in the reaction buffer Y (33 mM Tris-acetate (pH 7.9 @37C), 66 mM K-acetate) supplemented with 2 mM DTT at 37°C for 5 min. The reaction was started by the addition of 25 µM (unless otherwise stated) of cyclic or linear oligoadenylate. Samples were incubated at 37°C for 120 min (or other time point if specified otherwise) and the reaction was stopped by mixing with 4× SDS loading dye (200 mM Tris, 400 mM DTT, 8% (w/v) SDS, 40% (v/v) glycerol and 6 mM bromophenol blue). The reaction products were separated by SDS-PAGE (15% gel) and visualized by Coomassie staining. Gels were imaged using the densitometry scanning mode on an Amersham™ Typhoon™ Biomolecular Imager (Cytiva).

### cA_n_ synthesis reactions and cA_4_ radiolabeling

The mixture of 0.2 μM of CCaCsm complex, 10 μM of target RNA or non-specific RNA and 60 μM of ATP in the reaction buffer H (20 mM HEPES (pH 7.5), 50 mM KCl, 0.1 mg/ml BSA (Thermo Fisher Scientific) was supplemented with 1 mM of Mg(CH_3_COO)_2_ and the cA_n_ synthesis reaction was performed at 37°C for 75 min. The reaction was stopped by freezing the sample. The reaction products were subjected to HPLC-MS analysis. α^32^P-labeled cA_4_ for subsequent direct visualization of its hydrolysis was prepared by mixing 0.2 μM CCaCsm complex with 10 μM target RNA (5’-GGGUGAAGAGCAAUGAGCUCUCGAGGUGCGAUAUCGCUCUUCCCAGUGUA-3’

RNA prepared by *in vitro* transcription of annealed oligodeoxynucleotides as described in^67^), 1 mM Mg(CH_3_COO)_2_, 0.2 mM ATP and 1 μM α^32^P-ATP in the reaction buffer H and incubating at 37°C for 4 hours. Synthesis products were separated by denaturing PAGE (24% gel, 19:1 acrylamide:bisacrylamide ratio, 6 M urea in 0.5× TBE) and cA_4_ was purified from the gel by phenol extraction and ethanol precipitation.

### Ring nuclease reactions

The cA_n_ hydrolysis assay was performed in the reaction buffer Y supplemented with 1 mM DTT, 0.1 mg/mL BSA (Thermo Fisher Scientific), and 0.5 U/μl RiboLock RNase inhibitor (Thermo Fisher Scientific) containing 10 μM of synthetic cA_3_ (Biolog), synthetic cA_4_ (Biolog), synthetic cA_6_ (Biolog), linear A_4_>p (ChemGenes), or linear A_4_p (IDT). Reactions were initiated by adding 1 μM of protein (wt or mutant versions of CCaCalpL) and incubated at 37°C for the indicated time. Reactions were stopped by freezing the sample. For time course experiments, aliquots were taken at the indicated time points and the reactions were quenched by heating at 85°C for 10 min. The reaction products were subjected to HPLC-MS analysis.

To directly visualize the hydrolysis of cA_4_, reactions were performed in the reaction buffer supplemented with 1 mM DTT, 0.1 mg/mL BSA (Thermo Fisher Scientific), and 0.5 U/μl RiboLock RNase inhibitor (Thermo Fisher Scientific) and containing 50 nM CCaCsm-produced radiolabeled α^32^P-cA_4_. Multiple turnover reactions additionally contained 10 μM of unlabeled cA_4_ (Biolog). Reactions were started with the addition of 1 μM protein and run at 37°C. At the indicated time points, 5 μl aliquots were taken and the reactions were quenched by mixing with an equal amount of 2× RNA Loading Dye (Thermo Fisher Scientific) at 85°C. Reaction products were analyzed by denaturing PAGE (30% gels, 19:1 acrylamide:bis-acrylamide ratio, 6 M urea in 0.5× TBE) and visualized by autoradiography.

### HPLC-MS analysis

#### Analysis of cA_n_ synthesis and cA_4_ hydrolysis products

To analyze the cA_n_ synthesized by CCaCsm and the hydrolysis products of cA_n_, electrospray ionization mass spectrometry (ESI-MS) was performed in negative mode using an integrated HPLC/ESI-MS system (1290 Infinity, Agilent Technologies/Q-TOF 6520, Agilent Technologies) equipped with a Supelco Discovery®HS C18 column (7.5 cm × 2.1 mm, 3 µm). Elution was performed with a linear gradient of solvents A (5 mM ammonium acetate in water, pH 7.0) and B (5 mM ammonium acetate in methanol, pH 7.0) at a flow rate of 0.3 ml/min at 30°C as follows: 0-3 min, 0% B; 3-23 min, 15% B; 23-25 min, 40% B, 25-26 min 80% B, 26-29 min 80% B. Ionization capillary voltage was set to 5000 V, fragmentor to - 150 V. Molecules were annotated based on accurate mass, retention time of synthesized standards and fragment ions.

#### Analysis of CCaCalpT cleavage products

Samples were analyzed on an integrated HPLC/ESI-MS system (Agilent 1290 Infinity) equipped with a Poroshell 300SB-C8 column (2.1x75 mm, 5 µm) by elution with a linear gradient of solvents A (1% formic acid in water) and B (1% formic acid in acetonitrile) at a flow rate of 0.4 ml/min at 30°C as follows: 0-1 min, 2% B; 1-6 min, 2-98% B; 6-7 min, 98% B; 7-9 min, 98-2% B; 9-10 min, 2% B. High-resolution mass spectra of protein products were acquired on an Agilent Q-TOF 6520 mass analyzer (100-3200 m/z range, positive ionization mode). Results were analyzed using Agilent MassHunter Qualitative Analysis B.05.00 software.

### Cryo-EM sample preparation

The CCaCalpL complexes with oligoadenylates were prepared by mixing 10.5 µM (0.66 mg/ml) of N-tagged CCaCalpL with 30 µM of cA_4_, A_4_>p or A_4_p in the reaction buffer Y supplemented with 1 mM DTT and 2.5% glycerol just before applying on the freshly glow-discharged copper 300 mesh R1.2/1.3 holey carbon grids (Quantifoil), in a Vitrobot Mark IV (FEI) at 4 °C with a waiting time of 0 s and a blotting time of 5 s under 95% humidity conditions. Grids were plunge-frozen in liquid ethane cooled at liquid nitrogen temperature. A_4_>p and A_4_p samples were prepared on holey carbon grids overlaid with graphene oxide (GO). The GO coating was performed as described^68^.

### Cryo-EM data collection and image processing

The cryo-EM data for the CCaCalpL with oligoadenylate complexes were collected using a Glacios microscope (Thermo Fisher Scientific), running at 200 kV and equipped with a Falcon 3EC Direct Electron Detector in the electron counting mode (Vilnius University). Images were recorded with EPU (v.3.2) at a nominal magnification of ×92,000, corresponding to a calibrated pixel size of 1.10 Å per pixel, using an exposure of 0.80 e/Å^2^ s^−1^, in 30 frames and a final dose of 29.7 e/Å^2^, over a defocus range of −1.0 to −2.0 µm. Patch motion correction, CTF estimation, micrograph curation, blob picking, and particle extraction were performed in real-time in CryoSPARC Live (v.4.2.1)^69,70^. Further data processing was performed using standard CryoSPARC (v.4.2.1)^69,70^.

2,604,122 particles of the CCaCalpL-A_4_p complex were extracted (box size 240 pixels) from 3,127 accepted micrographs (Figure S4). After 2D classification, the selected particles (898,101) were subjected to heterogeneous refinement using four volumes obtained from an ab-initio reconstruction. Class 1 possessing higher FSC resolution (4.22 Å, 488,970 pct) was further subjected to non-uniform refinement followed by 3D classification into 3 classes. After 3D classification, particles from one class (175,828) were used for the final reconstruction using non-uniform refinement, local refinement, reference-based motion correction, followed by the final rounds of non-uniform and local refinement.

1,426,639 particles of the CCaCalpL-cA_4_ complex were extracted (box size 240 pixels) from 2,471 accepted micrographs. After 2D classification, the selected particles (687,353) were subjected to heterogeneous refinement using three volumes obtained from an ab-initio job. Class 2 possessing higher FSC resolution (4.22 Å, 339,216 pct) was further subjected to a second round of heterogeneous refinement into 3 classes. Particles from class 2 possessing higher FSC resolution (4.22 Å, 132,224 pct) were subjected to another round of 2D classification. Selected filament particles (124,470 pct) were used for the final reconstruction using non-uniform and local refinement.

3,472,781 particles of the CCaCalpL-A_4_>p complex were extracted (box size 240 pixels) from 3,127 accepted micrographs. After 2D classification, the selected particles (1,358,761 pct) were subjected to heterogeneous refinement using four volumes obtained from an ab-initio reconstruction. Class 1 possessing higher FSC resolution (4.22 Å, 559,559 pct) was further subjected to 3D classification to 3 classes. After 3D classification, particles from class 0 (215,062) were used for the final reconstruction using non-uniform and local refinement.

The global resolution and sphericity values for all reconstructions were estimated using 3DFSC v.3.0 software^71^ according to the Fourier shell correlation of 0.143 criterion^72^. The local resolution was estimated in CryoSPARC (v.4.3.0)^69,70^.

#### Cryo-EM model building and validation

The initial protein model for CCaCalpL-A_4_p complex was generated using AlphaFold^63^ under the ColabFold^64^ framework using default parameters and MMseqs2 to search for homologues into the ColabFold database, and manually modified using Coot (v.0.9.8.1)^73^ against the map sharpened using phenix.auto_sharpen (v.1.20.1-4487)^74^. A_4_p was built manually. Model refinement was performed using phenix.real_space_refine (v.1.20.1-4903)^74^. The final model of the CCaCalpL-A_4_p complex covers protein residues 4-13, 18-97, 104-166, 171-197, 201-505 of chain A; residues 4-30, 36-62, 66-96, 104-189, 201-422, 426-489, 492-506 of chain B; residues 203-351, 363-504 chain C; and two A_4_p molecules. This model was further used as the initial model for CCaCalpL-cA_4_ and CCaCalpL-A_4_>p complexes, which were rebuilt and refined using similar procedures. The final model of the CCaCalpL-cA_4_ complex covers protein residues 202-352, 357-504 of chain A; residues 202-229, 233-273, 279-337, 344-353, 356-422, 426-486, 496-506 of chain B a and one cA_4_ molecule. The final model of the CCaCalpL-A_4_>p complex covers protein residues 90-95, 107-112, 118-164, 174-188, 202-353, 357-505 of chain A; residues 154-167, 207-228, 231-337, 344-352, 358-419, 426-486, 494-504 of chain B and residues 220-300, 307-326, 333-350, 384-419, 422-473, 480-502 of chain C and two A_4_>p molecules.

The statistics of the three-dimensional reconstruction and model refinement are summarized in Table S2. The molecular graphics figures were prepared with ChimeraX (v.1.5)^75^.

### Biolayer Interferometry

Biolayer interferometry (BLI) experiments were performed using the Octet K2 system (ForteBio, Pall). Experiments were performed at 37°C in 96-well plate format using 200 μL reagent volumes and a stage rotation rate set of 1000 rpm.

#### CCaCalpL interaction with CCaCalpT-CalpS

The His_6_-StrepII-His_6_-tagged CCaCalpT-CalpS heterodimer was used as the ligand and the H396A or R358/K361E variants of the CCaCalpL protein as analyte. The Ni-NTA biosensors (ForteBio, U.K.) were hydrated in the reaction buffer Y supplemented with 1 mM DTT, 0.01% Triton X-100, 10 mM imidazole and 0.1% BSA (Merck) for 10 minutes before use. After the initial baseline step of 120 seconds, the CCaCalpT-CalpS heterodimer was immobilized on biosensors at a concentration of 200 nM for 120 seconds. A secondary baseline of 120 seconds was then performed to stabilize the signal. The 60-second association step was recorded by transferring the ligand-bound biosensors to wells containing the analyte CCaCalpL at concentrations of 250 nM, 500 nM, 1000 nM and 2000 nM. The association step was followed by a dissociation phase of 180 seconds. The reactions were supplemented with 4 μM of cA_4_ in the appropriate measurement series. A biosensor without CCaCalpT-CalpS ligand was used as a reference. Before each measurement in the series with increasing analyte concentration, the biosensors were regenerated by three cycles of incubation for 5 seconds in 10 mM glycine (pH 1.7), followed by 5 seconds in the kinetics buffer. The biosensors were then recharged by incubation in 10 mM NiCl_2_ for 60 seconds. The mean amplitude of the CCaCalpL binding response, normalized to the amplitude of the CCaCalpT-CalpS ligand loading response, is reported from three replicates for each CCaCalpL concentration.

#### CCaCalpS interaction with RNAP

The His_6_-StrepII-His_6_-tagged CCaCalpS, CCaCalpT-CalpS heterodimer, or CCaCalpT-CalpS heterodimer after cleavage with cA_4_-activated CCaCalpL, was used as the ligand and *E. coli* RNA polymerase core enzyme (New England Biolabs) as the analyte. The binding assay was performed in the 1× *E. coli* RNA Polymerase reaction buffer (New England Biolabs) supplemented with 0.01% Triton X-100, 10 mM imidazole and 0.1% BSA (Merck). Measurement steps were performed as described above for the CCaCalpL and CCaCalpT-CalpS binding experiment, except that the association step was extended to 90 seconds, and only a single concentration of RNAP (100 mU/µL) was used. We report the averaged BLI sensorgrams of three technical replicates for each immobilized protein.

### *E. coli* survivability assays

#### Spot assay

*E. coli* ER2566 cells were co-transformed with pCalpS or pCDFDuet-1 (Novogen) plasmid in combination with (i) pETDuet-1 (Novogen), (ii) pCalpT, (iii) pCalpT_24N_, (iv) pCalpT_11C_, or (v) pCalpT_split_ (Table S1), plated on LB media agar plates, supplemented with streptomycin (25 µg/ml), carbenicillin (50 µg/ml) and 1% glucose and grown overnight at 37°C. A single colony of each variant was grown overnight at 37°C in liquid LB supplemented with streptomycin (25 µg/ml), carbenicillin (50 µg/ml) and 1% glucose. The overnight cultures were serially diluted in fresh LB media and then plated onto LB media agar plates supplemented with streptomycin (25 µg/ml), carbenicillin (50 µg/ml) and 0.15 mM IPTG or 1% glucose. Bacteria were grown at 37°C for 16 hours. The experiment was performed in triplicate.

#### Time course experiment

Similar to the spot assay, *E. coli* ER2566 cells were co-transformed with the appropriate plasmids and the overnight cultures were obtained. The overnight cultures were diluted 1:200 in the fresh LB containing streptomycin (25 µg/ml) and carbenicillin (50 µg/ml) and incubated in a 96-well plate for 1 hour at 37°C with shaking. Then 0.1 mM IPTG was added to induce the protein expression. After induction bacterial growth was measured by recording OD_600_ every 10 minutes using the CLARIOstar Plus Microplate Reader. The mean with standard deviation of three biological replicates is presented for each experiment.

### Protein interaction assays using Dynabeads

#### CCaCalpS interaction with E. coli RNAP

In the reaction buffer (10 mM Tris-HCl (pH 7.5), 50 mM KCl, 1 mM DTT), 5 µM of the His_6_-StrepII-His_6_-tagged CCaCalpS was mixed with 400 mU/µL of *E. coli* RNA polymerase core enzyme (New England Biolabs) to a final volume of 30 µL. The proteins were allowed to associate by incubation at 37°C for 40 minutes. The reaction was diluted with reaction buffer H supplemented with 0.001% Triton X-100, 10 mM imidazole and 0.1 % BSA (Merck) to a final volume of 300 µL. The diluted reaction was then transferred to a tube containing 80 µg of pre-washed Dynabeads His-Tag Isolation and Pulldown magnetic beads (Invitrogen) and incubated on a roller for 5 minutes at room temperature. The supernatant was discarded and the beads were washed once with reaction buffer H supplemented with 0.001% Triton X-100, 10 mM imidazole and 0.1 % BSA (Merck), followed by three washes with the same buffer without BSA. Target proteins were eluted from the magnetic beads with 20 µL of reaction buffer H supplemented with 0.001% Triton X-100 and 300 mM imidazole and analyzed on a 12% SDS gel.

#### Protein interaction within CCaCalpT-CalpS heterodimer after protease cleavage

The protease reaction was performed by pre-incubating 5 µM of CCaCalpL and 5 uM of CCaCalpT-CalpS dimer in the reaction buffer Y supplemented with 2 mM DTT at 37°C for 5 minutes and starting the reaction by adding 25 µM of cA_4_. Samples were incubated at 37°C for 120 minutes in a final volume of 45 µL. The subsequent purification using magnetic beads was performed as described above for the CCaCalpS interaction with *E. coli* RNAP experiment, *except* reaction buffer Y with additives was used instead of reaction buffer H.

### GFP degradation in E. coli assay

#### Plate spot assay of GFP degradation

*E. coli* ER2566 cells were transformed with pGFP, pGFP-(wt)CalpT_24N_-degron, pGFP-(mut)CalpT_24N_-degron, or pGFP-ssrA and plated on LB medium agar plates, supplemented with carbenicillin (50 µg/ml) and grown overnight at 37°C. A single colony of each variant was grown overnight at 37°C in liquid LB supplemented with carbenicillin (50 µg/ml). The overnight cultures were serially diluted in fresh LB media and then plated onto LB media agar plates supplemented with carbenicillin (50 µg/ml) and 0.075 mM IPTG. Bacteria were grown at 37°C for 14 hours. Images of GFP-fluorescing cells on the agar plates were captured using the Amersham™ Typhoon™ Biomolecular Imager (Cytiva). Three biological replicates were performed.

#### Time-course measurements of GFP degradation

The overnight cultures (prepared as described above) were diluted 1:50 with fresh LB supplemented with ampicillin (50 µg/ml) and were grown to the mid-log phase. GFPmut1-degron expression was induced with 0.005 mM IPTG, and the cell suspension was further cultured overnight at 18°C. Bacterial cultures were diluted 1:10 in M9 minimal medium supplemented with 5% (w/v) glucose, 10 µg/mL thiamine, 10 µg/mL biotin, trace elements, and 100 µg/mL spectinomycin in a 96-well plate. Spectinomycin was used to stop protein translation. The OD_600_ and fluorescence (excitation at λ = 470 nm; emission at λ = 512 nM) were measured every 10 minutes for 12 hours at 30°C using a CLARIOstar Plus Microplate Reader. Fluorescence results were normalized first by optical density and then by setting the values at zero time points to 1. Results represent the mean with standard deviation of three biological replicates.

## QUANTIFICATION AND STATISTICAL ANALYSIS

Non-linear regression analysis of multipe-turnover cA_4_ cleavage reaction data was performed using Kyplot 2.0 software^76^. The reaction course was approximated by simultaneous fitting equations (1-4) that describe sequential conversion of cA_4_ to A_4_>p, A_4_p and A_2_>p + A_2_p over time with the rate constants k_1_, k_2_ and k_3_, respectively, to experimental data:

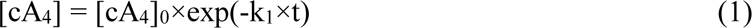

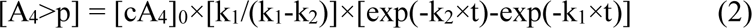

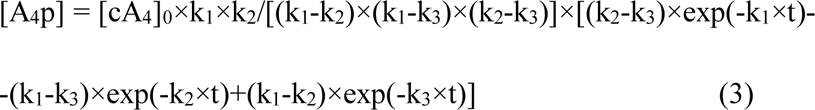

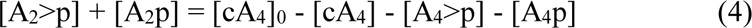

The observed turnover rates k_obs1_, k_obs2_ and k_obs3_ for the three reaction stages were then calculated by multiplying k_1_, k_2_ and k_3_ by the total substrate concentration [cA_4_]_0_ (10 µM) and dividing by the total enzyme concentration (1 µM) used in the reaction. The determined cA_4_, A_4_>p and A_4_p turnover rates are presented as the mean ± 1 standard deviation of k_obs1_, k_obs2_ and k_obs3_ values determined from three independent experiments.

## DATA AVAILABILITY

The cryo-EM 3D reconstructions have been deposited in the Electron Microscopy Data Bank under the accession codes EMD-50054 for CCaCalpL filament bound to A_4_p, EMD-50055 for CCaCalpL filament bound to cA_4_, EMD-50056 for CCaCalpL filament bound to A_4_>p. The cryo-EM model coordinates have been deposited in the Protein Data Bank under the accession code 9EYI (CCaCalpL filament bound to A_4_p), 9EYJ (CCaCalpL filament bound to cA_4_) and 9EYK (CCaCalpL filament bound to A_4_>p).

## ACKNOWLEDGMENTS

We thank Irmantas Mogila, Virginijus Siksnys and Česlovas Venclovas for the discussions. This work was supported by the Research Council of Lithuania (LMTLT) grant S-MIP-22-09 (grant to G. Tamulaitis).

## AUTHOR CONTRIBUTIONS

Conceptualization: D.S., G.Tamulaitiene, and G.Tamulaitis;

Methodology and Validation: D.S., and G.S.;

Formal analysis: D.S., and G.S;

Investigation: D.S., A.R., G.S., and G.Tamulaitiene;

Writing – original draft: D.S., G.S., G.Tamulaitiene, and G.Tamulaitis;

Writing – review and editing: D.S., G.S., G.Tamulaitiene, and G.Tamulaitis;

Visualization: D.S., G.S., and G.Tamulaitiene;

Supervision, Project administration and Funding acquisition: G.Tamulaitis.

