## Supplementary material for "Filament formation activates protease and ring nuclease activities of CRISPR SAVED-Lon": Figures S1-S8 and Tables S1-S2

**SUPPLEMENTAL FIGURES**

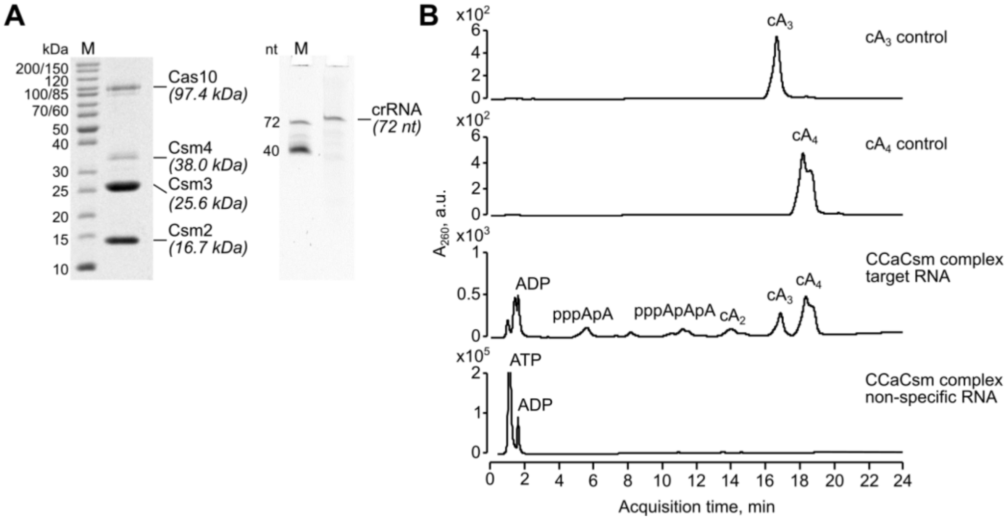


**Figure S1. $cA_4$ synthesis of *Candidatus* Cloacimonas acidaminovorans (CCa) Csm complex. Related to Figure 1.**

(A) Composition of the CCaCsm complex. Protein composition of the complex was analyzed by SDS-PAGE and Coomassie staining (left). M, protein molecular weight marker. Nucleic acid co-purifying with the complex was analyzed by denaturing PAGE and staining with SybrGold (right). M, DNA oligonucleotides of a known length, used as length markers.

(B) HPLC-MS analysis of $cA_n$ produced by CCaCsm complex in the presence and absence of target RNA. Synthetic $cA_3$ and $cA_4$ (Biolog) were used as controls.

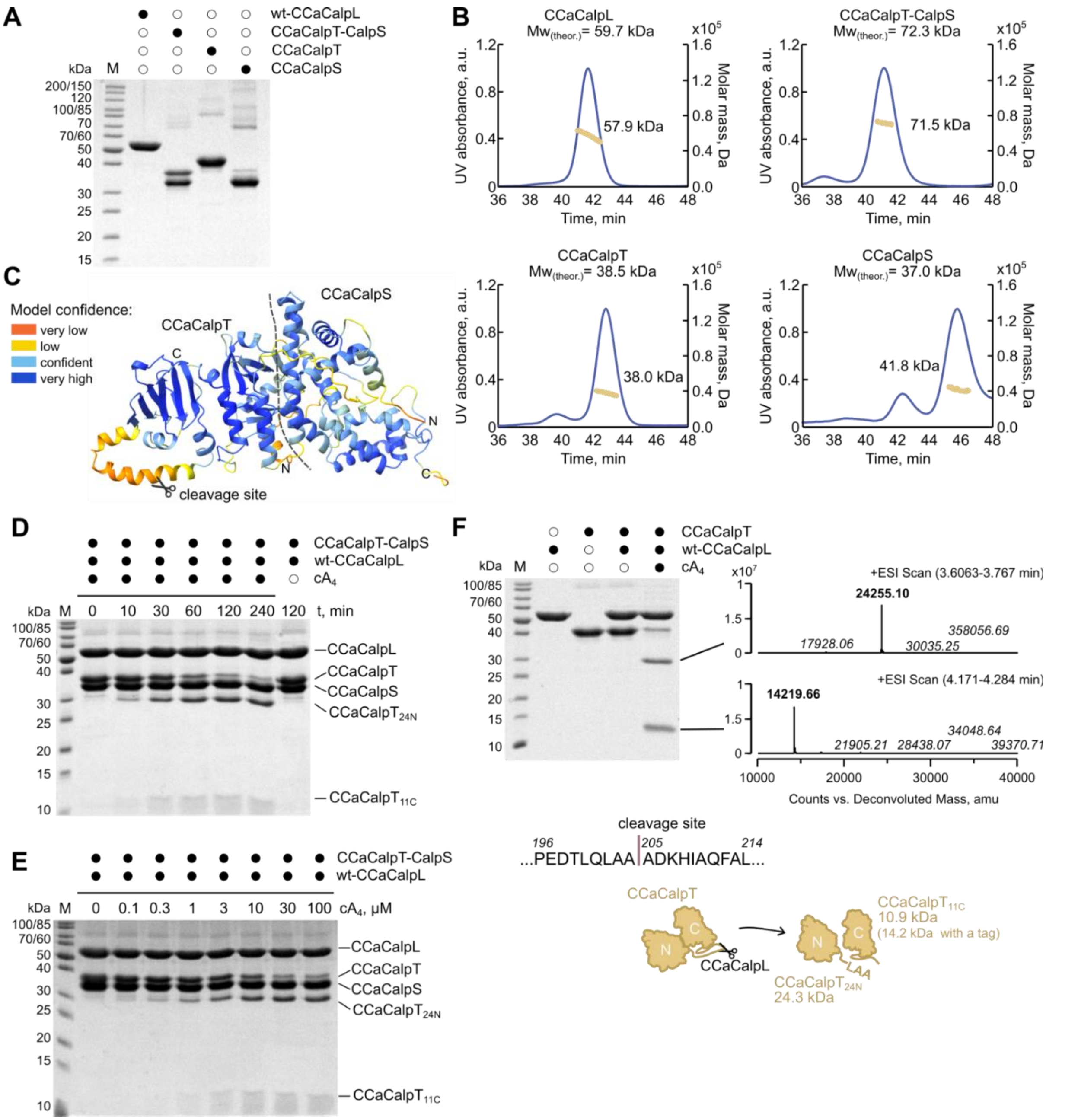


**Figure S2. Characterization of the tripartite CCaCalpL-CalpT-CalpS system. Related to Figures 1 and 4.**

(A) SDS-PAGE of purified wt-CCaCalpL, CCaCalpT-CalpS (CalpS C-tagged), CCaCalpT (C-tagged), CCaCalpS (C-tagged) proteins. M, protein molecular weight marker.

(B) SEC-MALS analysis of purified proteins: CCaCalpL (top left), CCaCalpT-CalpS (top right), CCaCalpT (bottom left), and CCaCalpS (bottom right). The calculated experimental molecular weights of CCaCalpL, CCaCalpT, CCaCalpT-CalpS correspond to protein monomer, heterodimer, monomer and monomer, respectively.

(C) AlphaFold2 models of CCaCalpT-CalpS heterodimer colored by model confidence score. Model confidence: very low (pLDDT <50); low (70> pLDDT >50); confident (90> pLDDT >70); very high (pLDDT >90).

(D) Time course experiment of CCaCalS-CalpT cleavage by $cA_4$-activated CCaCalpL. Aliquots were removed at timed intervals and samples were analyzed by SDS-PAGE and Coomassie staining. The reaction contained 5 µM CCaCalpT-CalpS heterodimer, 5 µM wt-CCaCalpL and 25 µM $cA_4$. Reaction without $cA_4$ served as a negative control. M, protein molecular weight marker.

(E) Dependence of CCaCalpL protease activity on activator $cA_4$ concentration. 120 min reactions contained 5 µM CCaCalpT-CalpS heterodimer, 5 µM wt-CCaCalpL and the indicated concentration of $cA_4$.

(F) Identification of the CCaCalpL cleavage site within the CCaCalpT target protein. The reaction contained 5 µM CCaCalpL, 5 µM CCaCalpT and 30 µM $cA_4$. Reaction products were analyzed by SDS-PAGE and Coomassie staining (top left) and by HPLC-MS (top right). Schematic depiction of CCaCalpT cleavage by CCaCalpL is shown at the bottom.

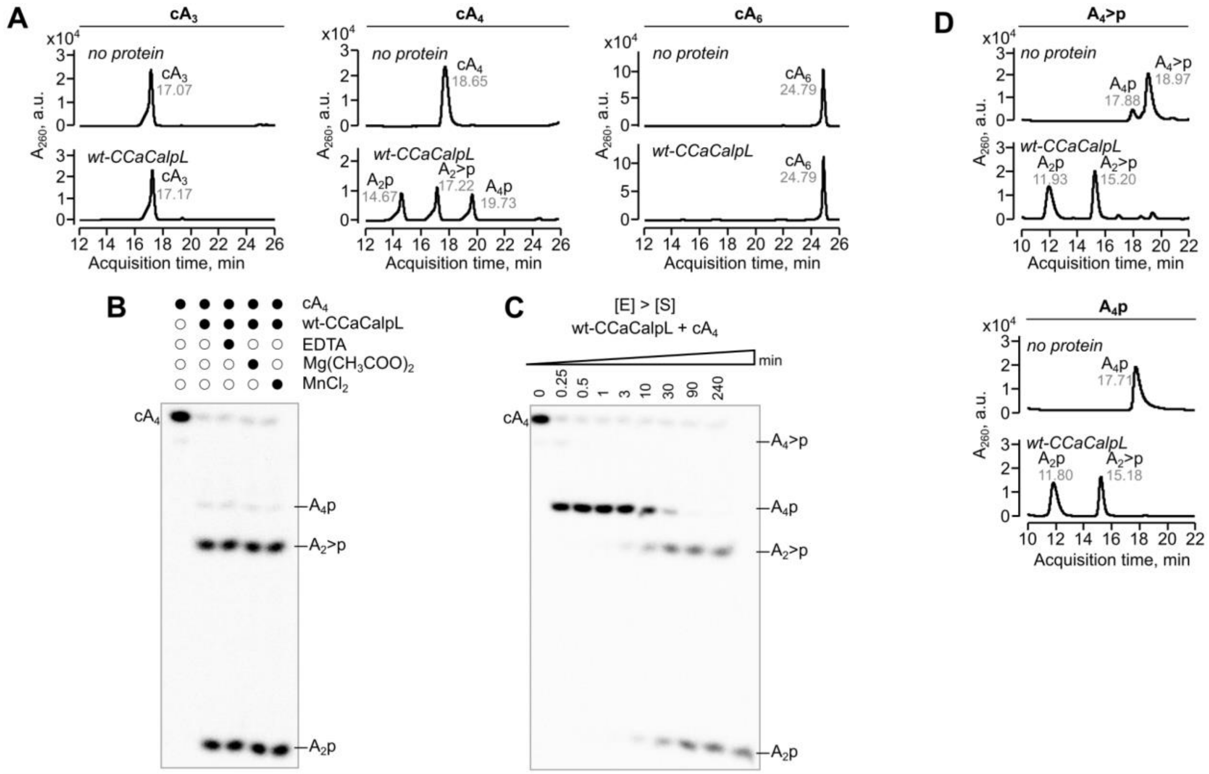


**Figure S3. CCaCalpL ring nuclease activity. Related to Figure 1.**

(A) Evaluation of wt-CCaCalpL ring nuclease specificity. wt-CCaCalpL cleavage reactions of $cA_3$, $cA_4$ and $cA_6$ were analyzed by HPLC-MS. The reactions contained 1 µM CCaCalpL and 10 µM $cA_n$.

(B) Cleavage of radioactively labeled $cA_4$ by wt-CCaCalpL in the presence or absence of divalent metal ions. 90 min reactions contained 1 µM CCaCalpL and 50 nM $\alpha^{32}$P-labeled $cA_4$ in the reaction buffer supplemented with either 1 mM EDTA, 10 mM Mg-acetate or 1 mM $MnCl_2$.

(C) CCaCalpL $cA_4$ cleavage reactions of radioactively labeled $cA_4$ by wt-CCaCalpL under single turnover conditions. Aliquots were removed at timed intervals and samples were analyzed by denaturing PAGE. The reaction contained 1 µM wt-CCaCalpL and 50 nM $\alpha^{32}$P-labeled $cA_4$.

(D) Cleavage of intermediates $A_4$>p and $A_4$p by wt-CCaCalpL. Cleavage products were analyzed by HPLC-MS.

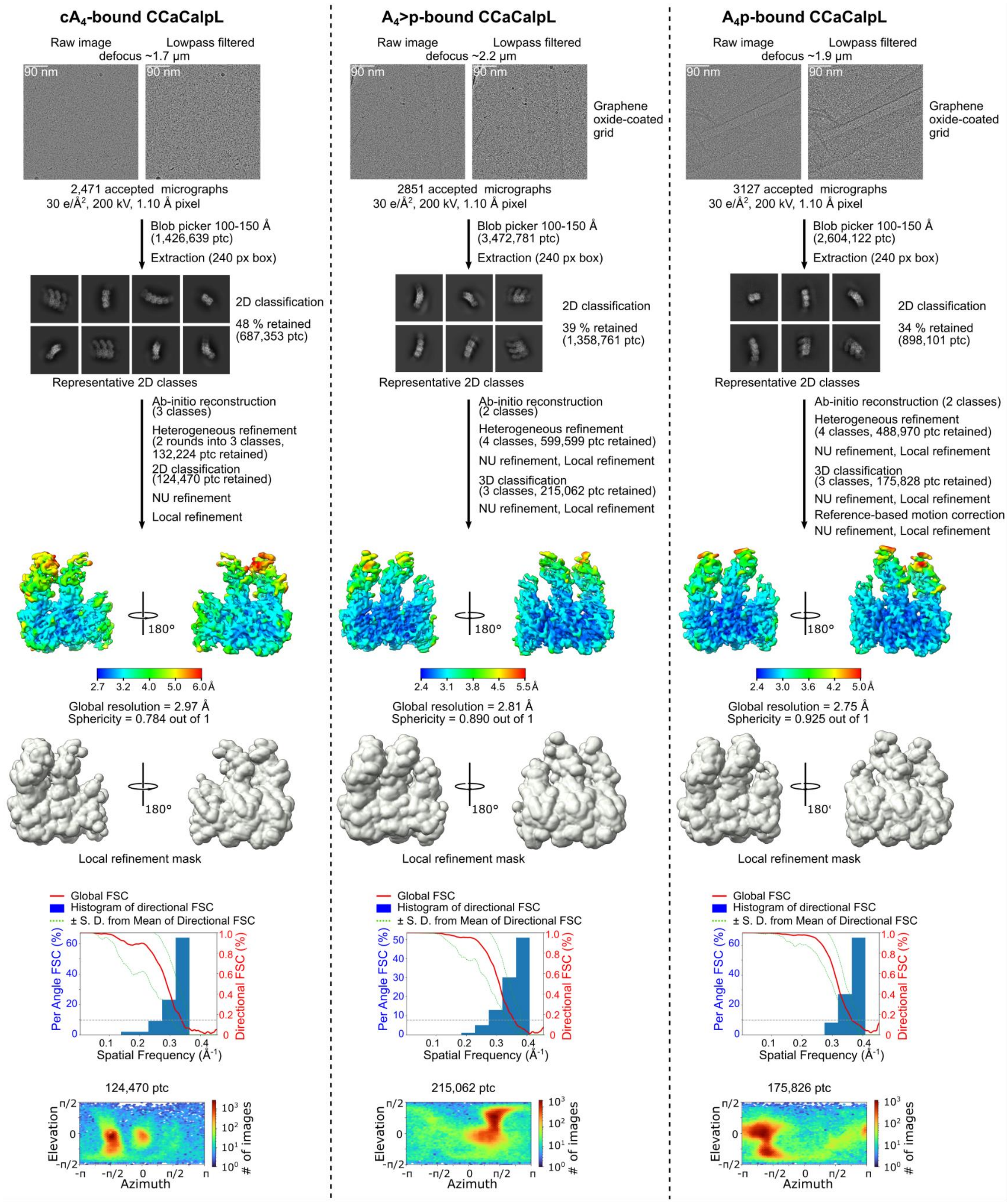


**Figure S4. Cryo-EM single particle analysis workflow. Related to Figure 2.**

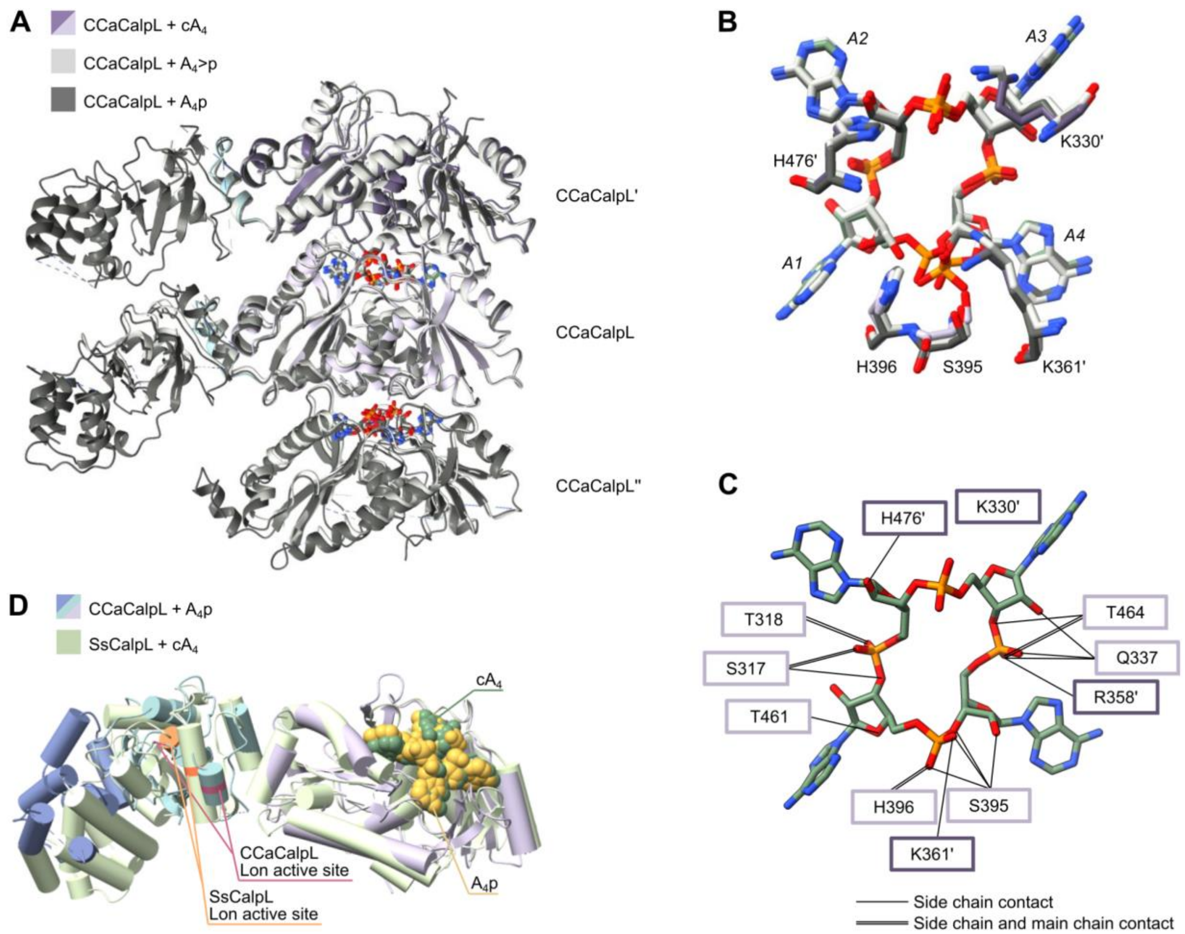


**Figure S5. Comparison of structures of CCaCalpL bound to $cA_4$, $A_4p$ and $A_4$>p. Related to Figure 2.**

(A) Superimposed structures of CCaCalpL bound to $cA_4$ (dark purple and light purple), $A_4$>p (light gray), and $A_4p$ (dark gray).

(B) The superposition of $cA_4$, $A_4$>p, and $A_4p$ bound to the CCaCalpL filament.

(C) CCaCalpL contacts with the phosphodiester backbone of $cA_4$. Single lines depict side chain contacts, double lines - side chain and main chain contacts. The residues from the CCaCalpL subunit are depicted in light purple squares, the residues of the adjacent CCaCalpL' subunit are shown in dark purple squares.

(D) The structure of a single monomer from the CCaCalpL filament resembles the structure of SsCalpL (PDB ID: 8B0R). The overlapping Lon active sites of both CalpLs are indicated.

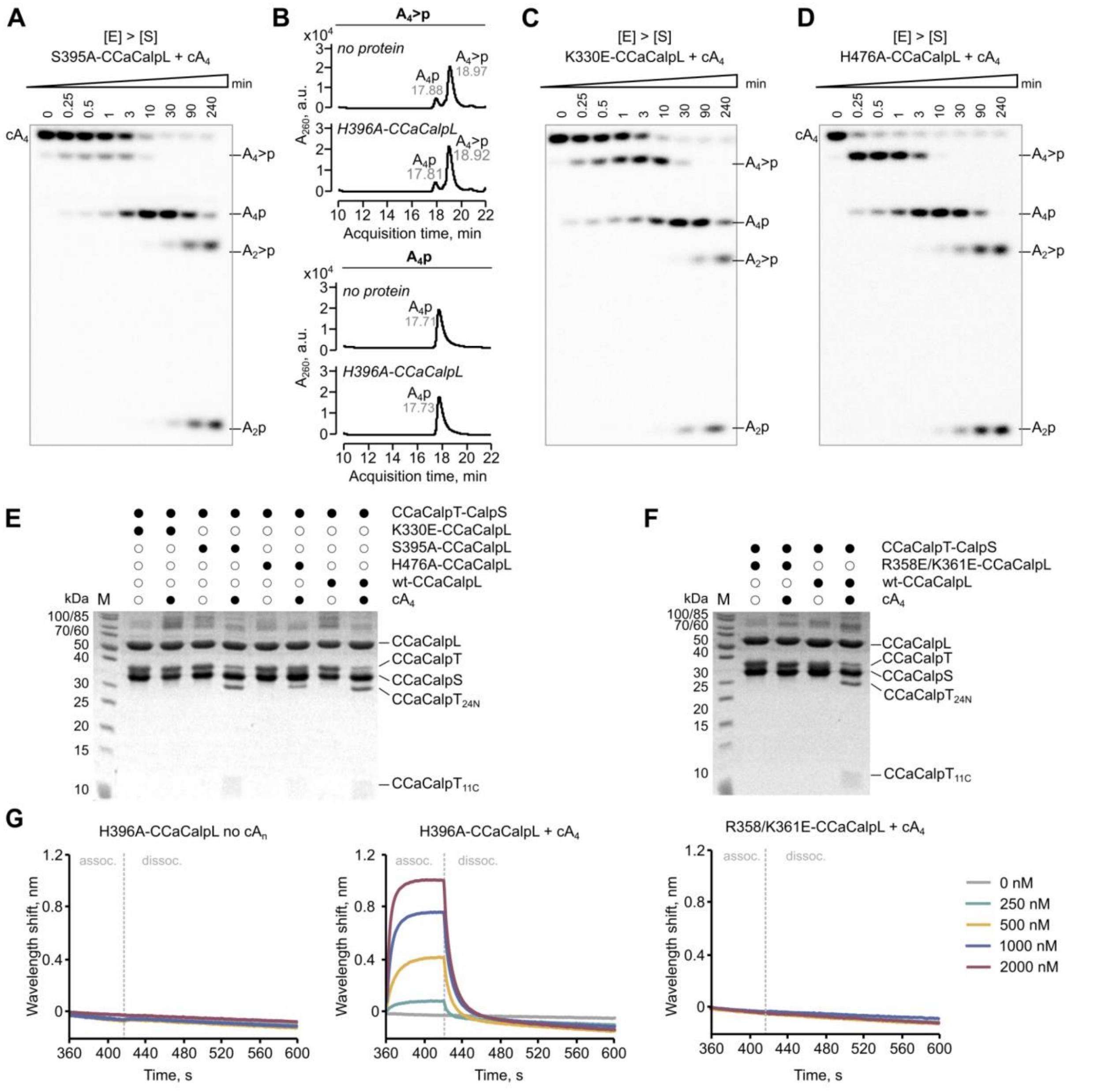


**Figure S6. CCaCalpL residues important for ring nuclease activity and filament formation. Related to Figures 2 and 3.**

(A) Cleavage of radioactively labeled $cA_4$ under single turnover conditions by S395A-CCaCalpL. Aliquots were removed at timed intervals and samples were analyzed by denaturing PAGE. The reactions contained 1 μM CCaCalpL and 50 nM $\alpha^{32}$P-labeled $cA_4$.

(B) Cleavage reactions of intermediates $A_4$>p and $A_4$p by H396A-CCaCalpL analyzed by HPLC-MS.

(C and D) Cleavage reactions of radioactively labeled $cA_4$ under single turnover conditions by K330E-CCaCalpL (C) and H476A-CCaCalpL (D). Reactions were performed and analyzed as in (A).

(E) Comparison between protease reactions of wt and ring nuclease mutant variants of CCaCalpL. The reactions contained 5 µM CCaCalpL, 5 µM CCaCalpT-CalpS and 25 µM $cA_4$. Reaction products were analyzed by SDS-PAGE and Coomassie staining. M, protein molecular weight marker.
(F) Comparison between protease reactions of wt-CCaCalpL and interface mutant R358E/K361E-CCaCalpL. Reactions were performed as in (E).
(G) Representative BLI sensorgrams showing H396A-CCaCalpL (in the presence or absence of 4 µM $cA_4$) and R358/K361E-CCaCalpL (in the presence of 4 µM $cA_4$) association and dissociation to CCaCalpT-CalpS immobilized on the biosensor.

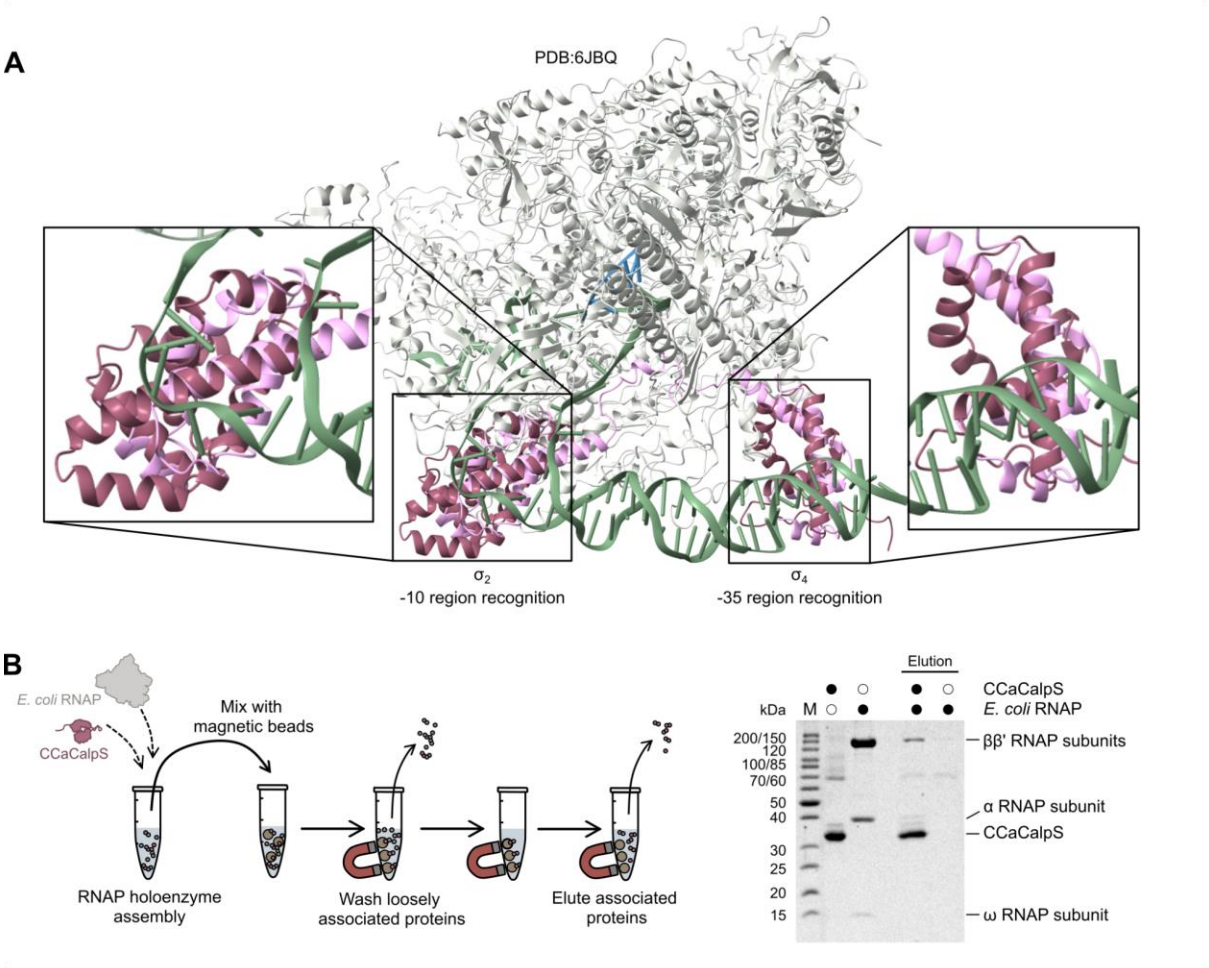


**Figure S7. CCaCalpS pull-down assay and toxicity studies in *E. coli*. Related to Figure 4.**
(A) AlphaFold model of CCaCalpS superimposed on *E. coli* RNAP/$\sigma^E$ transcription initiation complex (PDB: 6JBQ). RNAP is colored light gray, $\sigma^E$ – pink, DNA – green, RNA – blue, CCaCalpS – red. The linker region between the N-part and C-part of CCaCalpS has been cut to allow the superposition of the $\sigma^2$ and $\sigma^4$ domains, respectively, onto those of $\sigma^E$.
(B) CCaCalpS association with *E. coli* RNAP analyzed by pull-down assay. Schematic representation of the experiment (left). RNAP holoenzyme assembly was performed in the assembly buffer (10 mM Tris-HCl pH7.5, 50 mM KCl, 1 mM DTT) and contained 5 µM of C-tagged CCaCalpS and 400 mU/µl of *E. coli* RNAP core enzyme (NEB). The formed complex was pulled down using magnetic beads and analyzed by SDS-PAGE and Coomassie staining. M, protein molecular weight marker.

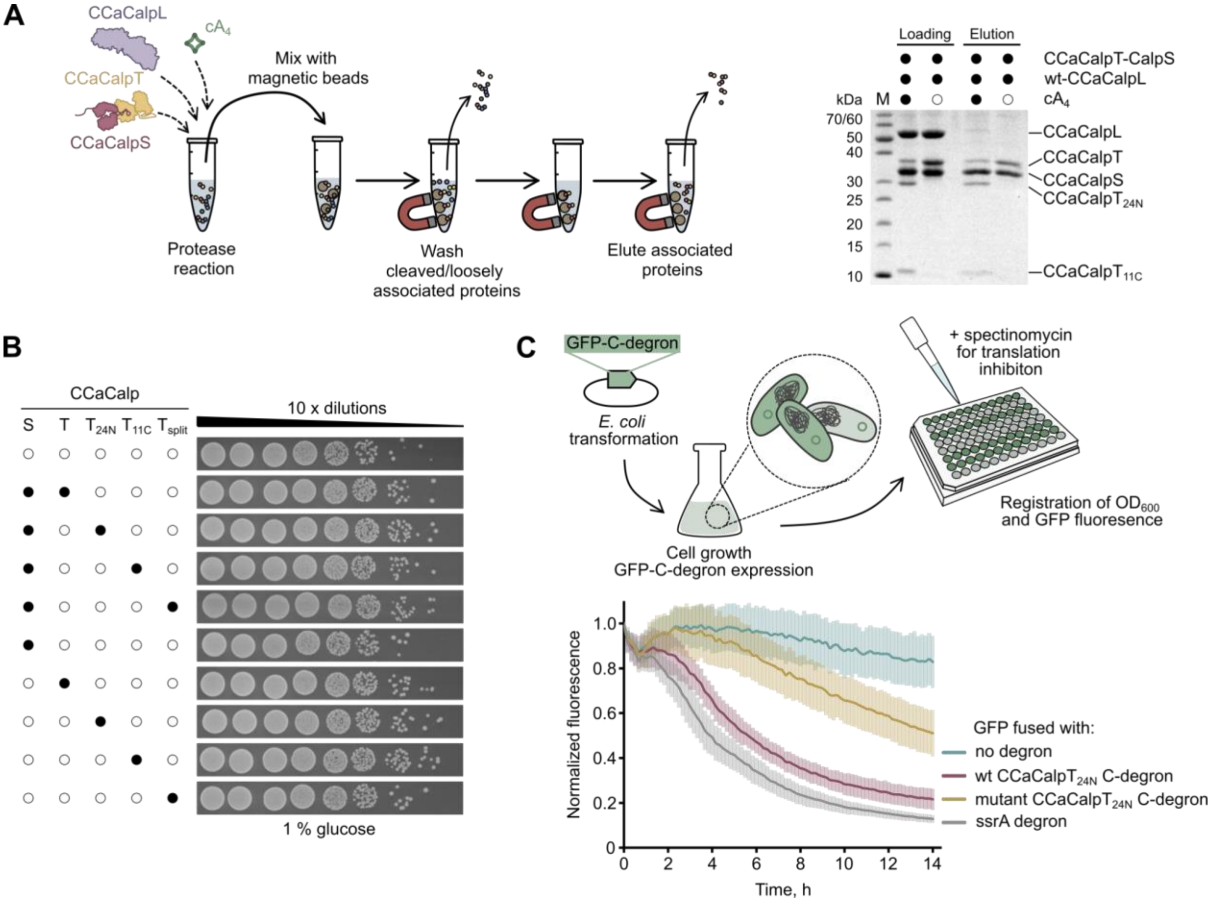


**Figure S8. Studies on the dissociation of CCaCalpS by pull-down assay and quantitative degradation analysis of GFP fused to various degrons. Related to Figure 4.**

(A) Testing the dissociation of CCaCalpS from CCaCalpT-CalpS heterodimer following CCaCalpL cleavage by pull-down assay. Schematic representation of the experiment is shown on the left. A protease reaction containing 5 µM wt-CCaCalpL, 5 µM CCaCalpT-CalpS heterodimer (with the affinity tag at the C-terminus of CCaCalpS) and either 25 µM $cA_4$ or no $cA_n$ was performed. The reaction mixture was then transferred to a tube containing $Ni^{2+}$-NTA-coated magnetic beads. Proteins loosely associated with CCaCalpS were washed out, and eluted CCaCalpS with tightly bound proteins was analyzed by SDS-PAGE (right).

(B) *E. coli* expressing combinations of CCaCalpS and CCaCalpT variants (full-length, $CalpT_{24N}$, $CalpT_{11C}$, or split) plated on repressive LB media supplemented with 1 % glucose. (C) Quantitative degradation assay of GFP fused with degrons (PETLQLAA - $CCaCalpT_{24N}$ C-degron, PETLQLAD - mutant version of $CCaCalpT_{24N}$ C-degron and ENYALAA - partial fragment of ssrA C-degron) in liquid *E. coli* cultures over time. Schematic representation of the experiment (top). Variants of GFP were expressed in the *E. coli*, the overnight cultures were diluted with M9 minimal media, and protein translation was stopped by adding spectinomycin. Fluorescence signal of GFP was normalized to optical density (bottom). All data points are presented as the mean values of three independent experiments ± 1 SD.

### SUPPLEMENTAL TABLES

#### Table S1. Plasmids used in this study. Related to Figures 1 and 4.

| Plasmid | Description | Usage |
|---|---|---|
| pCsm | pCDFDuet-1_Cas10$_{N\text{-}tag}$-Csm2-Csm3-Csm4-Csm5 (Str$^R$) | Plasmid for expression of protein subunits of *Ca.* C. acidaminovorans wt-Csm complex. |
| pCRISPR | pETDuet-1_(repeat-spacer-repeat)-Cas6 (Ap$^R$) | Plasmid for expression of *Ca.* C. acidaminovorans crRNA. Plasmid encodes *cas6* gene and minimal CRISPR region. |
| pCalpL-$^N$Tag | pBAD_CalpL$_{N\text{-}tag}$ (Ap$^R$) | Plasmid for expression of *Ca.* C. acidaminovorans wt-CalpL variant fused with a TEV-cleavable His$_6$-StrepII-His$_6$ tag at the N-terminus. |
| pCalpL-$^C$Tag | pBAD_CalpL$_{C\text{-}tag}$ (Ap$^R$) | Plasmid for expression of *Ca.* C. acidaminovorans wt-CalpL variant fused with a TEV-cleavable His$_6$-StrepII-His$_6$ tag at the C-terminus. |
| p(S154A)CalpL-Tag | pBAD_(S154A)CalpL$_{C\text{-}tag}$ (Ap$^R$) | Plasmid for expression of *Ca.* C. acidaminovorans S153A mutant CalpL variant fused with a TEV-cleavable His$_6$-StrepII-His$_6$ tag at the C-terminus. |
| p(S395A)CalpL-Tag | pBAD_(S395A)CalpL$_{C\text{-}tag}$ (ApR) | Plasmid for expression of Ca. C. acidaminovorans S394A mutant CalpL variant fused with a TEV-cleavable His$_6$-StrepII-His$_6$ tag at the C-terminus. |
| p(H396A)CalpL-Tag | pBAD_(H396A)CalpL$_{C\text{-}tag}$ (Ap$^R$) | Plasmid for expression of *Ca.* C. acidaminovorans H396A mutant CalpL variant fused with a TEV-cleavable His$_6$-StrepII-His$_6$ tag at the C-terminus. |
| p(K330E)CalpL-Tag | pBAD_(K330E)CalpL$_{C\text{-}tag}$ (Ap$^R$) | Plasmid for expression of *Ca.* C. acidaminovorans K330E mutant CalpL variant fused with a TEV-cleavable His$_6$-StrepII-His$_6$ tag at the C-terminus. |

| | | |
|---|---|---|
| p(R358E-K361E)CalpL-Tag | pBAD_(R358E-K361E)CalpL$_{C\text{-}tag}$ (Ap$^R$) | Plasmid for expression of *Ca.* C. acidaminovorans R358E and K361E mutant CalpL variant fused with a TEV-cleavable His$_6$-StrepII-His$_6$ tag at the C-terminus. |
| p(H476A)CalpL-Tag | pBAD_(H476A)CalpL$_{C\text{-}tag}$ (Ap$^R$) | Plasmid for expression of *Ca.* C. acidaminovorans H476A mutant CalpL variant fused with a TEV-cleavable His$_6$-StrepII-His$_6$ tag at the C-terminus. |
| pET-CalpS-Tag | pETDuet-1_CalpS$_{C\text{-}tag}$ (Ap$^R$) | Plasmid for expression of *Ca.* C. acidaminovorans wt-CalpS variant fused with a TEV-cleavable His$_6$-StrepII-His$_6$ tag at the C-terminus. Used in combination with pCDF-CCaCalpT for CCaCalpT-CalpS co-expression. |
| pCDF-CalpT | pCDFDuet-1_CalpT (Str$^R$) | Plasmid for expression of *Ca.* C. acidaminovorans wt-CalpT variant without a tag. Used in combination with pET-CalpS-C for CalpT-CalpS co-expression. |
| pET-CalpT-Tag | pETDuet-1_CalpT$_{C\text{-}tag}$ (Ap$^R$) | Plasmid for expression of *Ca.* C. acidaminovorans wt-CalpT variant fused with a His$_6$-StrepII-His$_6$ tag at the C-terminus. |
| pCalpS | pCDFDuet-1_CalpS (Str$^R$) | Plasmid encoding *Ca.* C. acidaminovorans *calpS* for the toxicity assay in *E. coli*. |
| pCalpT | pETDuet-1_CalpT(1-299) (Ap$^R$) | Plasmid encoding *Ca.* C. acidaminovorans full-length (1-299 codons) *calpT* for the toxicity assay in *E. coli*. |
| pCalpT$_{24N}$ | pETDuet-1_CalpT(1-204) (Ap$^R$) | Plasmid encoding *Ca.* C. acidaminovorans *calpT$_{24N}$* (1-204 codons) for the toxicity assay in *E. coli*. |
| pCalpT$_{11C}$ | pETDuet-1_CalpT(205-299) (Ap$^R$) | Plasmid encoding *Ca.* C. acidaminovorans *calpT$_{11C}$* (205-299 codons) for the toxicity |

| | | |
|---|---|---|
| | | assay in *E. coli*. |
| pCalpT$_{split}$ | pETDuet-1_CalpT(1-204)_CalpT(205-299) (Ap$^R$) | Plasmid encoding *Ca.* C. acidaminovorans *calpT$_{split}$* (1-204 and 205-299 codons) for the toxicity assay in *E. coli*. |
| pCalpT$_{24N}$(mut) | pETDuet-1_(A204D)CalpT(1-204) (Ap$^R$) | Plasmid encoding *Ca.* C. acidaminovorans *calpT$_{24N}$* (1-204 codons) with the A204D mutation in degron sequence for the toxicity assay in *E. coli*. |
| pGFP | pETDuet-1_GFP | Plasmid encoding GFP. Negative control for GFP degradation assays. |
| pGFP-(wt)CalpT$_{24N}$-degron | pETDuet-1_GFP-PETLQLAA | Plasmid encoding GFP at the C-terminus fused with CalpT$_{24N}$ C-degron PETLQLAA for GFP degradation assays. |
| pGFP-(mut)CalpT$_{24N}$-degron | pETDuet-1_GFP-PETLQLAD | Plasmid encoding GFP at the C-terminus fused with a mutant version of CalpT$_{24N}$ C-degron PETLQLAD for GFP degradation assays. |
| pGFP-ssrA-degron | pETDuet-1_GFP-ENYALAA | Plasmid encoding GFP at the C-terminus fused with ssrA C-degron ENYALAA. Positive control for GFP degradation assays. |
| pCDFDuet-1 | pCDFDuet-1 (Str$^R$) | Control plasmid used in the toxicity assay in *E. coli*. |
| pETDuet-1 | pETDuet-1 (Ap$^R$) | Control plasmid in the toxicity assay in *E. coli*. |

**Table S2. Cryo-EM data collection, refinement and validation statistics.**

**Related to Figure 2.**

| PDB<br>EMDB | **CCaCalpL-cA$_4$**<br>**9EYJ**<br>**EMD-50055** | **CCaCalpL-A$_4$>p**<br>**9EYK**<br>**EMD-50056** | **CCaCalpL-A$_4$p**<br>**9EYI**<br>**EMD-50054** |
|---|---|---|---|
| **Data collection and processing** | | | |
| Microscope | Glacios (TFS) | Glacios (TFS) | Glacios (TFS) |
| Detector | Falcon III camera (TFS) | Falcon III camera (TFS) | Falcon III camera (TFS) |
| Magnification | 92,000 | 92,000 | 92,000 |
| Automation software | EPU | EPU | EPU |
| Voltage (kV) | 200 | 200 | 200 |
| Electron exposure (e-/Å$^2$) | 29.7 | 29.7 | 29.7 |
| Defocus range (μm) | 0.5 to 2.5 | 0.5 to 2.5 | 0.5 to 2.5 |
| Pixel size (Å) | 1.1 | 1.1 | 1.1 |
| Symmetry imposed | C1 | C1 | C1 |
| Micrographs collected/used | 2,546/2,471 | 3,017/2,851 | 3,469/3,127 |
| Initial/final particle images (no.) | 1,426,639/124,470 | 3,472,781/215,062 | 2,604,122/175,826 |
| Map resolution (Å) (masked/unmasked); FSC threshold 0.143 | 2.97/3.18 | 2.81/2.94 | 2.75/2.92 |
| Map sharpening B factor (Å$^2$) | 105.8 | 108.5 | 119.7 |
| 3D FSC sphericity score | 0.784 | 0.890 | 0.925 |
| **Refinement** (Phenix) | | | |
| Initial model used | CCaCalpL-A$_4$p | CCaCalpL-A$_4$p | AlphaFold |
| Model resolution (Å) FSC threshold 0.5 | 3.38 | 3.29 | 3.00 |

| | | | |
|---|---|---|---|
| Model composition<br>Non-hydrogen atoms<br>Protein residues<br>Nucleotides | <br>4334<br>576<br>4 | <br>6639<br>891<br>8 | <br>9453<br>1248<br>8 |
| B factors (Å$^2$)<br>Protein | 72.07 | 80.57 | 83.48 |
| Nucleotide | 51.65 | 63.93 | 54.42 |
| RMSDs<br>Bond lengths (Å)<br>Bond angles (°) | <br>0.004<br>0.587 | <br>0.005<br>0.750 | <br>0.005<br>0.633 |
| **Validation** | | | |
| MolProbity score | 2.22 | 2.28 | 1.99 |
| Clashscore | 13.32 | 13.61 | 10.21 |
| Poor rotamers (%) | 2.14 | 3.10 | 2.15 |
| Ramachandran plot<br>Favored (%)<br>Allowed (%)<br>Disallowed (%) | <br>95.16<br>4.84<br>0 | <br>96.13<br>3.87<br>0 | <br>96.72<br>3.28<br>0 |
| CCvolume/CCmask | 0.71/0.72 | 0.73/0.73 | 0.81/0.81 |
| Cb outliers (%) | 0.18 | 0 | 0 |
| CaBLAM outliers (%) | 0.93 | 0.98 | 0.42 |
